# Geographical based variations in white truffle *Tuber magnatum* truffle aroma is explained by quantitative differences in key volatile compounds

**DOI:** 10.1101/2020.09.30.321133

**Authors:** Jun Niimi, Aurélie Deveau, Richard Splivallo

## Abstract

- The factors that vary the aroma of *Tuber magnatum* fruiting bodies are poorly understood. The study determined the headspace aroma composition, sensory aroma profiles, maturity, and microbiome composition from *T. magnatum* originating from Italy, Croatia, Hungary, and Serbia, and tested if truffle aroma is dependent on provenance and if fruiting body volatiles are explained by maturity and/or microbiome composition.
- Headspace volatile profiles were determined by gas chromatography-mass spectrometry-olfactometry (GC-MS-O) and aroma of fruiting body extracts were sensorially assessed. Fruiting body maturity were estimated through spore melanisation. Bacterial community was determined using 16S rRNA amplicon sequencing.
- Main odour active compounds were present in all truffles but varied in concentration. Aroma of truffle extracts were sensorially discriminated by sites. However, volatile profiles of individual fruiting bodies varied more within sites than across geographic area, while maturity level did not play a role. Microbiome composition varied highly and was partially explained by provenance. A few rare bacterial operational taxonomical units associated with select few non-odour active volatile compounds.
- Specificities of the aroma of *T. magnatum* truffles are more likely linked to individual properties than provenance. Some constituents of the microbiome may provide biomarkers of provenance and be linked to non-odour active volatiles.

## Introduction

The luxury standing of truffle fungi makes them one of the most expensive food in the world. Hundreds of truffle species exist, yet the white truffle *Tuber magnatum* is anecdotally regarded by many as the best and certainly as the most expensive truffle in the world. Retail Depending on the season coupled with low supply and high demand, prices range from 3,000-5,000 € per kg and can even reach as high as 7,000 € per kg (Riccioni *et al*., 2016).

*T. magnatum* is commonly known as the Alba or the Piedmont truffle since it occurs naturally in the Italian region of Piedmont, where truffles were already well known in the late middle ages (Rittersma, 2011). Yet, the natural distribution of *T. magnatum* is not limited to the region of Piedmont in northern Italy, but extends to most of the Italian territory, south east of France and the eastern European countries of Hungary, Croatia (Istria), Serbia, Bulgaria, Greece, Slovenia, and Romania (Marjanović *et al*., 2015; Belfiori *et al*., 2020). The reasons why *T. magnatum* grows only in these particular countries remain uncertain, but it is possible that these areas have suitable ecological conditions such as soil chemical composition and porosity with access to water for the growth and development of *T. magnatum* ectomycorrhizas and fruiting bodies (Bragato & Marjanović, 2016). Recent genetic studies on 36 *T. magnatum* populations encompassing more than 400 truffles have exemplified a clear genetic structure within Europe. Specifically, the presence of four genetic clusters for *T. magnatum* located in (i) southern Italy, (ii) central Italy and Istria, (iii) northern Italy, and (iv) the Balkan/Pannonian regions were revealed (Rubini *et al*., 2005; Belfiori *et al*., 2020). The localisation of these four genetic clusters further supports the inference that *T. magnatum* might have recolonized Europe after the last ice age starting from central Italy (Belfiori *et al*., 2020).

Truffles owe their reputation in big part to their unique and powerful smell. More than 60 volatile compounds were detected in *T. magnatum* but the smell of the white truffle can only be attributed to a little more than 10 molecules that are perceivable through human olfaction (Schmidberger & Schieberle, 2017). At its core is 2,4-dithiapentane, a sulfur containing compound that has a characteristic garlic and sulphur aroma of *T. magnatum*. This molecule was isolated and characterized from *T. magnatum* already in the 70’s (Fiecchi *et al*., 1967) and its synthetic version has since widely been used in truffle flavoured food products (Wernig *et al*., 2018). Other important contributors to *T. magnatum* aroma include 3-(methylthio)propanal (potato character), 2- and 3-methylbutanal (malty character) and 2,3-butanedione (buttery character) (Schmidberger & Schieberle, 2017). It is possible to recombine 11 synthetic volatile compounds in specific proportions and thereby reproduce the overall aroma character of *T. magnatum* (Schmidberger & Schieberle, 2017).

A recent study on *T. magnatum* listed 115 volatiles emitted by its fruiting bodies (Vita *et al*., 2018), which is a factor of magnitude higher than the odour active compounds reported previously (Schmidberger & Schieberle, 2017). Vita *et al*. (2018) further reported the presence of site specific volatile compounds within truffles from among 16 locations in Italy and one site in Croatia (Istria), suggesting volatile markers that exclusively occurred in specific regions, confirming earlier results by Gioacchini *et al*. (2008). Many of these site markers were sulfur-containing compounds or terpenes, two classes of compounds that are well known for their sulfurous, floral or citrus-like odours. Site markers might hence influence truffle aroma, provided that the markers are odour active and at levels above perception threshold. Recently, volatile marker compounds that can classify *T. magnatum* truffles based on country of origin were preliminarily reported, where markers for Slovenian truffles are ethanol, benzaldehyde, 2-methyl-1-butanol, and dimethyl sulfide, and those for Italian truffles are anisol, 1,4-dimethoxy-benzene, 1-methoxy-3-methyl-benzene, 1-octen-3-ol, and 2-methyl-butanal (Strojnik *et al*., 2020). Whether these compounds are exclusive to specific countries or that they were ubiquitous but present in varying proportions by country is unknown. Further, whether these compounds contribute to perceptual differences in aroma and if additional marker compounds from multiple countries exist are also unknown.

From the aforementioned studies, volatiles measured from truffles appear to vary across geographical origin, but the role of specific factors that might vary across provenance which in turn potentially influence white truffle aroma remain mysterious/debated. Numerous factors have been suggested to influence truffle aroma including fruiting body maturity (Zeppa *et al*., 2004), possibly the association with the host trees (Gioacchini *et al*., 2008; Vita *et al*., 2018), genetics (Splivallo *et al*., 2011; Molinier *et al*., 2015; Vahdatzadeh & Splivallo, 2018), microbes colonizing truffle fruiting bodies (Splivallo *et al*., 2015; Vahdatzadeh *et al*., 2015), and other environmental factors (i.e. soil, climate). Closely related to environmental factors is seasonal variation. As with other plants and fungi, it is possible that each of those factors could influence truffle aroma to a certain extent, yet, there is currently no consensus. References given above would suggest for instance that maturation might have an influence on the aroma of *Tuber borchii* but not on *Tuber aestivum*. This question has not yet been raised for *T. magnatum*.

The objective of this study was to shed light on some of the open questions on *T. magnatum* raised above. Applying techniques in aroma analysis, sensory science and microbial ecology, *T. magantum* truffles originating from seven orchards and four countries were analysed. First, the identity of volatile compounds and hence the aroma of truffles that may be behind potential differences were investigated. Second, the bacterial microbiome of individual truffle fruiting bodies from the varying provenances were determined. Thirdly, fruiting body maturation, weight, truffle provenance and microbiome were analysed as potential factors that could explain aroma variability among single truffle fruiting bodies.

## Materials and Methods

### Chemicals

Silicon oil, 2,3-butanedione (99 %), dimethyl sulfide (DMS) (≥ 99 %), (E)-2-octenal (95 %), trans-2-hexenal (96 %), isovaleric acid (99 %) were purchased from VWR (Darmstadt, Germany). 2,3-pentanedione (≥ 96 %), dimethyl sulfone (98 %), 1-octen-3-ol (98 %), 2-methyl butanal (95 %), 3-methyl butanal (97 %), heptanal (≥ 95 %), hexanal (98 %), (E,E)-2,4-nonadienal (≥ 89 %), methional (≥ 97 %), benzaldehyde (≥ 98 %), benzeneacetaldehyde (≥ 95 %), 2,4-dithiapentane (DTP) (≥ 99 %), dimethyl disulfide (DMDS) (≥ 99 %), dimethyl trisulfide (DMTS) (≥ 99 %), nonanal (≥ 95 %), 2-methyl-2-pyrroline, and alkane series standard solution (C8-C20) in hexane were purchased from Sigma-Aldrich (Taufkirchen, Germany). Short chain alkane standard mixture (C5-C8) was prepared in house; pentane, hexane, heptane (99 %) (VWR), and octane (99 %) (Sigma-Aldrich).

### Biological material and sample processing

Truffle fruiting bodies of *T. magnatum* were collected from natural truffle orchards in four countries (Italy, Hungary Serbia and Croatia) during one truffle season (October 2018 and January 2019 -Table 1). Two locations were sampled per country apart from Croatia, where truffles were collected from a single location, and at least four truffles were collected per site/truffle orchard (Table 1). More precise locations are not provided due to confidentiality requirements by the truffle-hunters who provided the samples. Once harvested, truffles were wrapped in paper towels, cooled to 4°C and transported/shipped to the laboratory within 3 days, where they were further processed as explained hereafter. Species identification was confirmed for every single fruiting body by spore morphology (when visible) and by PCR using *T. magnatum* species-specific primers as published earlier (Rizzello *et al*., 2012).

**Table 1.** Sample information of the fruiting bodies investigated.

| **Location**<br>**(region)** | **Sample**<br>**code** | **Collection**<br>**date** | **# fruiting**<br>**bodies** | **Mean maturity**<br>**(% ± SE)** | **Mean weight**<br>**(g ± SE)** |
| --- | --- | --- | --- | --- | --- |
| Hungary 1<br>(Baranya) | HUN1 | 10/2018 | 7 | 81.8 ± 2.0 | 11.8 ± 1.2 |
| Hungary 2<br>(Somogy) | HUN2 | 11/2018 | 4 | 74.7 ± 2.6 | 29.6 ± 6.6 |
| Italy 1*<br>(Abruzzo) | ITA1 | 11/2018 | 6 | 67.1 ± 5.9 | 31.3 ± 13.9 |
| Italy 2**<br>(Abruzzo) | ITA2 | 11/2018 | 5 | 68.8 ± 2.3 | 39.9 ± 3.5 |
| Serbia 1<br>(Kalubara) | SER1 | 12/2018 | 6 | 83.9 ± 1.4 | 24.8 ± 2.6 |
| Serbia 2<br>(Srem) | SER2 | 12/2018 | 5 | 90.6 ± 1.7 | 30.4 ± 3.3 |
| Croatia 1<br>(Istria) | CRO1 | 01/2019 | 7 | 45.1 ± 5.0 | 28.3 ± 7.2 |

\* and \*\*: ITA1 and ITA2 correspond to two natural but distinct truffle orchards located in Abruzzo region

The following processing steps, summarized in Fig. 1 that highlights the experimental design, were undertaken for samples upon arrival to the laboratory. Excess dirt was removed from each truffle with a brush under running cold water and truffles were dried with paper towels. The mass of each cleaned fruiting body was recorded before removing the outer layer (peridium) with sterile knives. For each fruiting body, multiple subsamples of fruiting body gleba were taken to analyse: maturity (75 ±25 mg) (Methods S1) (Fig. **1** – **2a**), volatiles (300 ±5 mg) (Methods S1) (Fig. **1** – **2b**), and the microbiome characterisation (75 ±25 mg) (Methods S3) (Fig. **1** – **2c**). The maturity and microbiome subsamples were placed in sterile 1.5ml Eppendorf tubes and kept at −20°C until further processing. The subsample for volatile analysis on the other hand, were analysed immediately after processing and measured fresh. We validated earlier that a single truffle subsample was sufficient to properly represent the volatilome of each truffle (Splivallo *et al*., 2012). The remainder of the gleba of the fruiting bodies from a single site were grated with a kitchen appliance electric grater (WMF, Geislingen) to yield a pooled sample of truffle representing a single site. The grated truffles from a given site were mixed with a sterile spoon before being sampled into 11 solid phase microextraction (SPME) vials (300 ±5 mg) in preparation for olfactory trials (total 8 vials per sample) (Fig. **1 - 1a**) and volatile analysis (total of 3 vials per sample) (Fig. **1 - 1b**). The volatile analysis and the determination of odour active compounds was performed using the gas chromatography-mass spectrometry (GC-MS) coupled with olfactometry (GC-O) (for further details, see Methods S2). For these analyses, grated fruiting bodies in SPME vials were held at 5°C overnight until olfactometry sessions the following day. The remainder of the grated fruiting bodies were combined with silicon oil at a ratio of 1:2 (50 g truffle: 100 g silicon oil). Silicon oil was chosen as the extraction media of choice through benchtop testing, where the oil was the most neutral in aroma characteristic and its ability to extract aroma of truffles that were the most similar to the fruiting bodies themselves. This mixture of grated fruiting bodies and silicon oil was homogenised (Ultra-Turrax®, IKA, Staufen, Germany), followed by centrifugation in 30ml centrifuge tubes at 8,000g for 10 min at 5°C (Heraeus Megafuge 8R centrifuge, Thermo Fisher Scientific, Osterode am Harz, Germany).

**Fig. 1.**
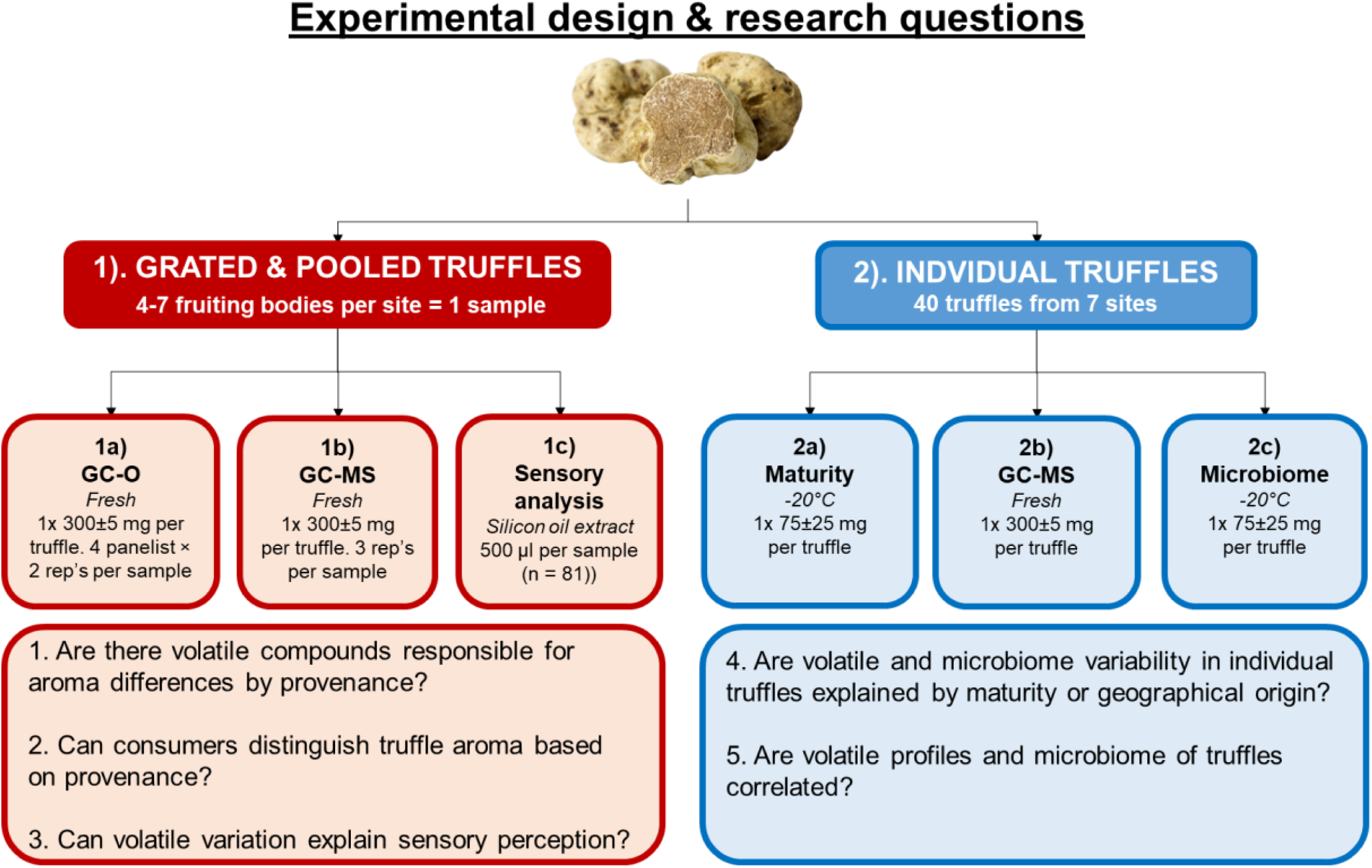
Experimental design for the analysis of individual truffles and pooled truffles and key questions associated with each design.

The silicon oil supernatant was decanted from the centrifuge tubes into 40ml amber vials with Teflon lined caps and stored at −20°C until required for sensory evaluation using the rate all that apply (RATA) methodology (Fig. **1 - 1c**). For a detailed description of the RATA, refer to Methods S4 and Table S1. The consumers who participated in the study were 57% female and 43% male. Most of the participants belonged to 18 −24 and 25 – 34 age groups (45.7% each), while that of 35 −44 and 55 −64 were minor (6% and 3%, respectively). The majority of consumers (74%, n = 81), were familiar with or had previous exposure to truffle products, while the remainder were unfamiliar.

### Data Analysis

The volatile data obtained from GC-MS analysis were imported as CDF files, and deconvoluted with Paradise (ver. 3, Copenhagen) (Johnsen *et al*., 2017). The resultant extracted compounds were integrated and calculated for their retention index. Compound identification was performed by comparing the electron ionisation mass spectra of the measured compounds either with the NIST 2017 GC-MS database (NIST, MD, USA), pure standards, or to literature for mass spec fragmentation patterns where required. Retention indices (RI) for the obtained features were calculated based on an alkane series (C5-C8 and C8-C20) (van Den Dool & Kratz, 1963). The peaks of chromatograms from individual fruiting bodies were normalised by dividing individual peaks by the total ion count and analysed with principle component analysis (PCA) using Unscrambler (ver. 10, Camo, Oslo). The peaks of chromatograms from the pooled truffle fruiting bodies (measured as with the GC-O samples) were processed in a similar manner as above, only that the odour active compounds identified (based on GC-O) were quantified through calibration curves. The quantified odour active compounds were subsequently analysed using one-way ANOVA using SPSS statistics Ver. 25 (SPSS, Inc., Chicago) and significantly different means were analysed with Fisher’s least significant difference (LSD) posthoc test.

GC-O data were manually aligned across all assessors for both replicates per sample. Where compounds were not detected by certain panellists, data were replaced with 0 scores. The entire data matrix was analysed using univariate ANOVA taking samples and panellist as fixed and random effects, respectively.

Data from RATA (sensory evaluation) were first preprocessed by replacing missing values with 0 scores. The preprocessed data was analysed using univariate ANOVA, taking samples and consumers as fixed and random effects, respectively. Post hoc testing was further performed on significantly different attribute means using Tukey’s Honest Significant Difference (HSD) test.

Details of bioinformatic processing of sequencing data are given in Supporting Information Methods S3. In brief, obtained sequences from amplicon sequencing were analysed using FROGS (Find Rapidly OTU with Galaxy Solution) (Escudié *et al*., 2017) following Standard Operation Procedures. Rare OTUs (≤ 0.005 % of all sequences in all samples) were excluded for further analyses. Clusters were affiliated to one taxonomy by blasting OTUs against SILVA database (Quast *et al*., 2012). OTUs were rarefied (adjusting sequences randomly to the total abundance in the smallest sample) to 45,846 using Phyloseq package in R (McMurdie & Holmes, 2013). The raw data are deposited in the NCBI Sequence Read Archive website (http://www.ncbi.nlm.nih.gov/sra) under the BioProject study accession number PRJNA663751.

The volatile profiles from pooled truffle samples were correlated with the RATA data using multiple factor analysis (MFA) (XLSTAT ver. 2020, Addinsoft, New York). The volatile profiles were taken as the mean concentration of compounds identified in the pooled truffle samples and the sensory data was taken as the significantly different (*P* < 0.05) attribute means. The correlation of the data matrices was further reported with RV coefficients.

The volatile profiles from individual truffle samples were correlated with the microbiome data using regularised canonical correlation analysis (rCCA) through the mixOmics R package (Rohart *et al*., 2017). Volatile profiles of individual fruiting bodies were correlated against the microbiome data of individual fruiting bodies. Data input were volatile profiles normalised to the total ion count within each fruiting body and rarefied reads of OTUs that were found in at least three fruiting body.

## Results

### Truffle odorants are ubiquitous among sites but vary in concentrations

The headspace of fresh truffle fruiting bodies, grated and pooled together by sites (Fig 1-2a) were determined for odour active compounds by olfactometry (GC-O) and a total of 25 odour active compounds were detected by olfaction across all samples (Fig. **2**). All compounds (apart from hexanal, 1,2,4-trithiolane, and compound RI1106) were ubiquitous across all samples, indicating that there were very little unique compounds present within a single site. However, a total of 48% of compounds significantly differed in perceived odour intensities among the sample sites (*P <* 0.05, Fig. **2a**). Yet, the significantly different intensities of compounds were based on individual sites rather than by country of origin. The intensities of the thirteen other compounds were similar across the samples, suggesting their ubiquity in the general composition of truffle aroma profile (Fig **2b**). The three compounds that did not appear to be ubiquitous from an olfaction point of view does not exclude the possibility that all three were present in all samples but for some at levels below olfactory detection threshold. Further, none of them were unique to any single site or region.

**Fig. 2.**
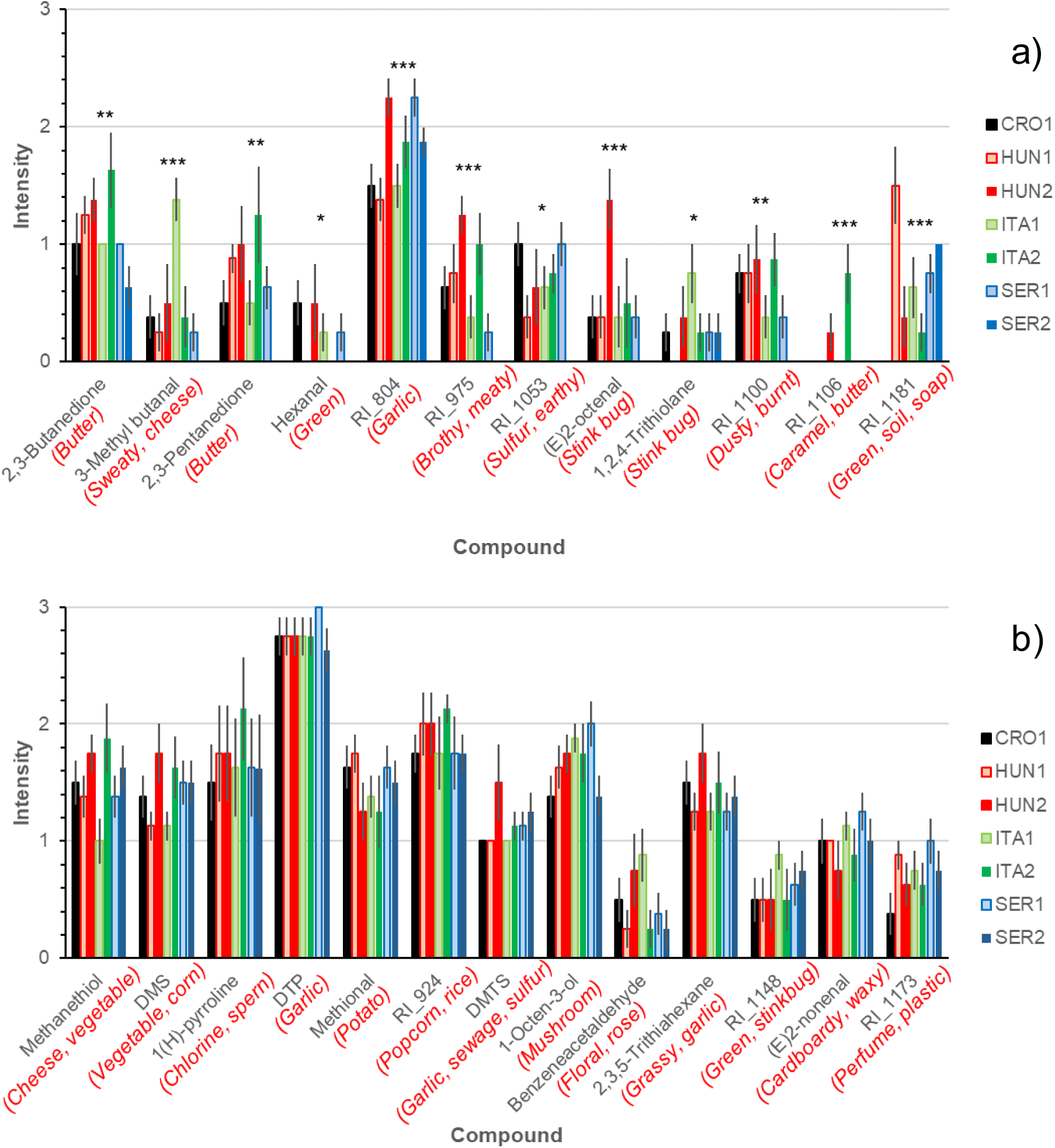
Mean odour intensities (± SE) measured from GC-O of ubiquitous compounds identified with accompanying odour descriptions in parentheses (n = 4 panellists). a) compounds that significantly differed in intensity across sample sites and b) compounds that did not significantly differ across samples. Compounds with numbers are retention indices (RI_) calculated through HP5-MS. Statistics according to univariate ANOVA; *, *P* < 0.05; **, *P* < 0.01; ***, *P* < 0.001.

Concentrations of volatile compounds detected in the headspace of pooled and grated truffle samples were determined for 13 odour active compounds that were quantifiable (Fig **1**-**2b**), of which 12 significantly differed (*P <* 0.05) in concentration across sites (Table 2.). For instance, ITA1 had the highest concentration of 3-methyl butanal consistently with the GC-O. The characteristic compound of *T*. magnatum, DTP, was detected in general at high concentrations, with SER2 containing the highest concentration. Interestingly, the odour intensity of DTP from GC-O were very high across all sites and were not significantly different. This is likely because the scale resolution could not capture very high perception intensities. There were compounds that were significantly different in concentrations but did not of their perceived intensities across sites, such as methanethiol, DMS, (1H)-pyrolline, DMTS, 1-octen-3-ol, and benzeneacetaldehyde. In these cases, it is likely that the difference in concentrations detected did not exceed perceived difference thresholds for each of these compounds, i.e. the minimum concentration difference for a compound to be perceivably different. Finally, concentrations of 2,3,5-trithiahexane were not significantly different between the sites. This was reflected on the high intensities detected by the assessors through the olfactometry in all samples (Fig. **2**). The combination of GC-MS of pooled truffle fruiting body data together with GC-O showed that both data are necessary for a more accurate inference in possible variations in aroma profiles of *T. magnatum*.

**Table 2.** Mean concentrations (μg kg) (± SE) of odour active volatile compounds from the headspace of seven fruiting body samples, pooled together as sites.

| Sample | RI* | CRO1 | HUN1 | HUN2 | ITA1 | ITA2 | SER1 | SER2 | ID | P-value |
| --- | --- | --- | --- | --- | --- | --- | --- | --- | --- | --- |
| Methanethiol<sup>§</sup> | <500 | 1,949<sup>b</sup><br>(± 169) | 1,373<sup>a</sup><br>(± 279) | 1,390<sup>ab</sup><br>(± 206) | 936<sup>a</sup><br>(± 136) | 1,408<sup>ab</sup><br>(± 45) | 2,403<sup>b</sup><br>(± 207) | 5,223<sup>b</sup><br>(± 191) | MS, Std,<br>RI, O | <0.001 |
| DMS | 531 | 25,279<sup>a</sup><br>(± 11,992) | 104,618<sup>ab</sup><br>(± 48,838) | 187,153<sup>b</sup><br>(± 19,644) | 62,706<sup>a</sup><br>(± 34,672) | 38,373<sup>a</sup><br>(± 9,501) | 96,921<sup>ab</sup><br>(± 4,934) | 164,046<sup>b</sup><br>(± 39,202) | MS, Std,<br>RI, O | 0.020 |
| 2,3-Butanedione | 600 | 10.0<sup>b</sup><br>(± 0.2) | 7.0<sup>b</sup><br>(± 1.4) | 8.8<sup>b</sup><br>(± 2.6) | 1.7<sup>a</sup><br>(± 0.6) | 4.4<sup>ab</sup><br>(± 0.6) | 4.5<sup>ab</sup><br>(± 1.0) | 3.7<sup>ab</sup><br>(± 0.2) | MS, Std,<br>RI, O | 0.005 |
| 3-Methylbutanal | 653 | 290<sup>a</sup><br>(± 13) | 381<sup>a</sup><br>(± 13) | 383<sup>a</sup><br>(± 42) | 1,082<sup>b</sup><br>(± 94) | 334<sup>a</sup><br>(± 16) | 395<sup>a</sup><br>(± 58) | 390<sup>a</sup><br>(± 34) | MS, Std,<br>RI, O | <0.001 |
| (1H)-Pyrroline<sup>†</sup> | 685 | 96<sup>b</sup><br>(± 10) | 56<sup>ab</sup><br>(± 1) | 44<sup>a</sup><br>(± 0) | 74<sup>b</sup><br>(± 13) | 44<sup>a</sup><br>(± 2) | 68<sup>ab</sup><br>(± 22) | 66<sup>ab</sup><br>(± 10) | MS<sup>†</sup>,<br>RI, O | 0.013 |
| 2,3-Pentanedione | 701 | 28<sup>b</sup><br>(± 2) | 29<sup>b</sup><br>(± 3) | 43<sup>c</sup><br>(± 4) | 14<sup>a</sup><br>(± 2) | 50<sup>c</sup><br>(± 6) | 13<sup>a</sup><br>(± 3) | 12<sup>a</sup><br>(± 3) | MS, Std,<br>RI, O | <0.001 |
| Hexanal | 802 | 366<sup>d</sup><br>(± 9) | 46<sup>ab</sup><br>(± 2) | 168<sup>c</sup><br>(± 87) | 344<sup>d</sup><br>(± 24) | 29<sup>a</sup><br>(± 4) | 395<sup>d</sup><br>(± 31) | 117<sup>bc</sup><br>(± 8) | MS, Std,<br>RI, O | <0.001 |
| DTP | 892 | 19,256<sup>a</sup><br>(± 1,467) | 24,102<sup>a</sup><br>(± 8,367) | 32,197<sup>a</sup><br>(± 2,938) | 19,995<sup>a</sup><br>(± 4,668) | 17,751<sup>a</sup><br>(± 1,206) | 38,423<sup>a</sup><br>(± 1,160) | 147,327<sup>b</sup><br>(± 17,773) | MS, Std,<br>RI, O | <0.001 |
| DMTS | 968 | 274<sup>a</sup><br>(± 4) | 277<sup>a</sup><br>(± 2) | 397<sup>b</sup><br>(± 65) | 279<sup>a</sup><br>(± 7) | 271<sup>a</sup><br>(± 1) | 290<sup>a</sup><br>(± 5) | 288<sup>a</sup><br>(± 3) | MS, Std,<br>RI, O | 0.037 |
| 1-Octen-3-ol | 980 | 4.33<sup>c</sup><br>(± 0.07) | 2.42<sup>a</sup><br>(± 0.04) | 3.36<sup>b</sup><br>(± 0.43) | 3.34<sup>b</sup><br>(± 0.03) | 2.21<sup>a</sup><br>(± 0.02) | 5.34<sup>d</sup><br>(± 0.42) | 3.80<sup>bc</sup><br>(± 0.13) | MS, Std,<br>RI, O | <0.001 |
| Benzeneacetaldehyde | 1045 | 1.62<sup>a</sup><br>(± 0.06) | 4.44<sup>b</sup><br>(± 0.11) | 5.89<sup>b</sup><br>(± 2.06) | 4.70<sup>b</sup><br>(± 0.51) | 1.84<sup>a</sup><br>(± 0.38) | 4.57<sup>b</sup><br>(± 0.52) | 5.97<sup>b</sup><br>(± 0.21) | MS, Std,<br>RI, O | 0.005 |
| (E)2-Octenal | 1057 | 5.73<sup>c</sup><br>(± 0.06) | 4.82<sup>ab</sup><br>(± 0.02) | 5.35<sup>bc</sup><br>(± 0.42) | 5.54<sup>c</sup><br>(± 0.18) | 4.70<sup>a</sup><br>(± 0.002) | 5.89<sup>c</sup><br>(± 0.14) | 5.35<sup>bc</sup><br>(± 0.12) | MS, Std,<br>RI, O | 0.004 |
| 2,3,5-Trithiahexane<sup>ψ</sup> | 1124 | 48<br>(± 4) | 47<br>(± 11) | 211<br>(± 88) | 93<br>(± 30) | 31<br>(± 1) | 99<br>(± 10) | 99<br>(± 14) | MS, RI,<br>O | 0.059 |

<sup>§</sup>Quantified as equivalent of dimethyl sulfide; <sup>†</sup>Quantified as equivalent of 2-methyl-1-pyrroline; <sup>ψ</sup>Quantified as equivalent of 2,4-dithiapentane; <sup>†</sup>MS fragmentation pattern based on literature (Schmidberger & Schieberle, 2017); \* RI calculated using HP5-MS. Statistical analysis was performed using one-way ANOVA and means with the same superscript are not significantly different according to Fishers LSD.

### Consumers can discriminate representative aroma of T. magnatum by site

In a second step, we tested whether consumers could distinguish truffles depending on their geographical origin by evaluating the aroma attributes (determined from the GC-O test) of fruiting bodies pooled by sites. Six attributes significantly differed (univariate ANOVA; *P <* 0.05) from a total of 14 attributes presented (Fig. **3**). The samples HUN2, and SER2 were significantly more intense in garlic aroma than the other samples, according to the panel of 81 consumers. CRO1 was characterised by earthy and chlorine character, while HUN1 was significantly more intense in potato and popcorn. ITA2 showed the highest intensity in cabbage aroma, while ITA1 was more intense in popcorn, though the overall range of intensities for popcorn was low. The sample SER1 did not discriminate strongly by any attribute. The data demonstrated here that even with extracts of truffle fruiting bodies, the consumers could discriminate the different samples qualitatively and quantitatively. Among these six attributes, the provenance of truffles could be discerned by site at a global level.

**Fig. 3.**
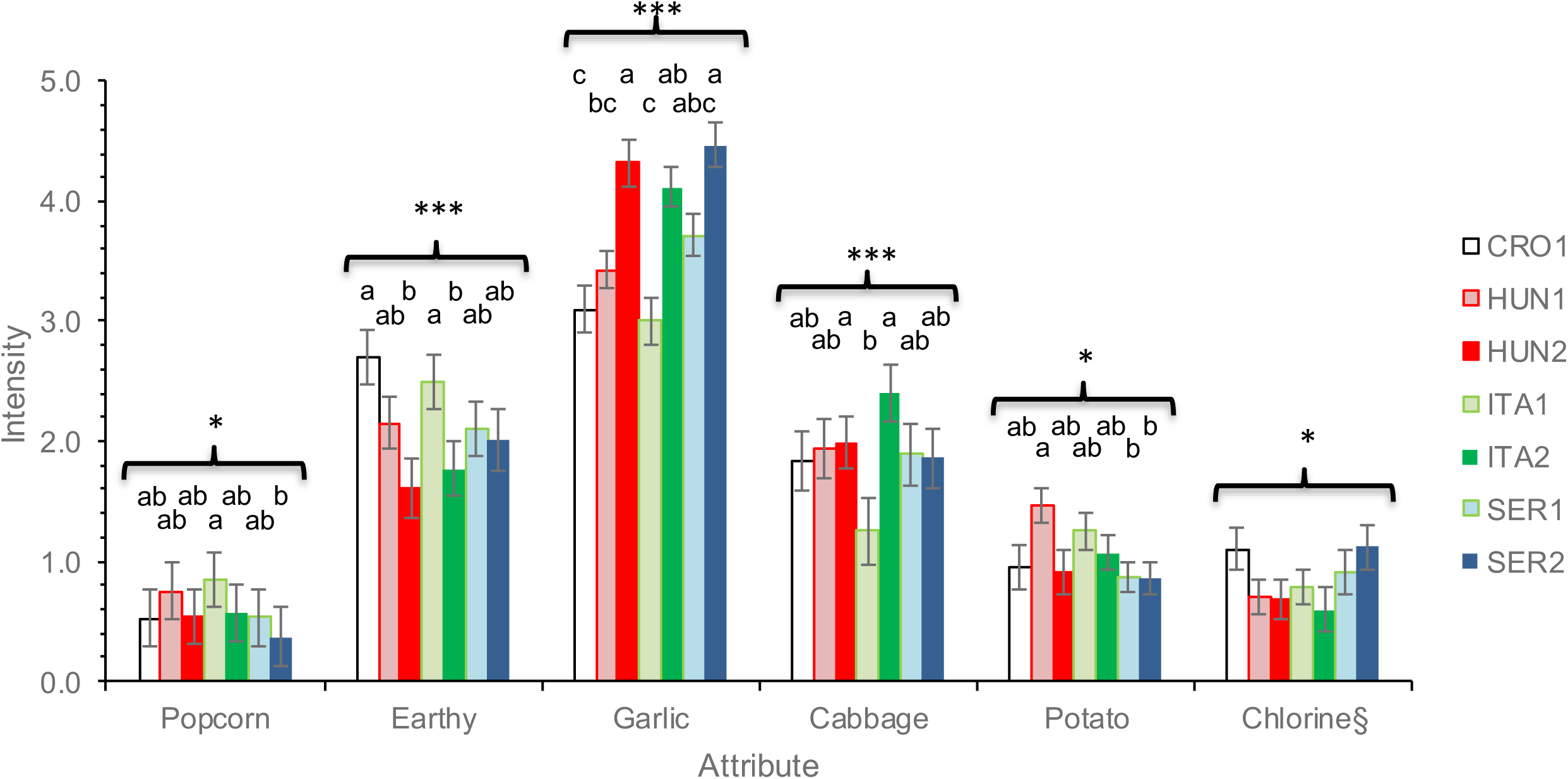
Mean intensities (± SEM) of significantly different RATA attributes (n = 81). ITA = Italy, HUN = Hungary, SER = Serbia, and CRO = Croatia. Statistical analysis is according to univariate ANOVA; *, *P* < 0.05; **, *P* < 0.01; ***, *P* < 0.001. Means with the same superscript per attribute are not significantly different according to Tukey’s HSD post hoc test. §Despite “chlorine” being a significantly different attribute by sample, Tukey’s HSD could not separate the means.

### Correlation between the volatile profiles and the sensory characteristics perceived by consumers

Single volatile compounds often have an odour character but can contribute differently to sensory characteristics when they are present in mixtures with other volatile compounds. To determine the compounds responsible for the perceived differences in global truffle aromas across sites, the quantified volatile data (GC-MS of pooled fruiting bodies) were correlated with the data obtained from sensory analysis (RATA) using MFA (Fig. **4**). Sensory attributes were overall projected with similar vectors as the compounds responsible, such as garlic with 2,3,5-trithiahexane, DTP, and DMTS, chlorine with (1H)-pyrroline (Fig. **4a**). 3-methylbutanal individually as a compound was perceived as sweaty and cheese from GC-O, yet when assessed in mixtures, it may have contributed to a global perception of earthy, popcorn, or potato. Likewise, the sulphur compounds may have had a role in eliciting the cabbage characteristic, perceived by the RATA panellists. Nevertheless, volatiles (GC-MS) and sensory data (RATA) were in agreement with each other (Fig. **4b**), indicating that both methods provide a good proxy of aroma variability at the pooled truffle fruiting body level. The largest disparity was seen with ITA2 sample, where this was mainly driven by a combination of comparatively higher concentrations of 2,3-pentanedione and lower concentrations of hexanal, DTP, and 2,3,5-trithiahexane (Fig. **4c**). The projected scores from volatiles and sensory data of the remaining samples were closely aligned.

**Fig. 4.**
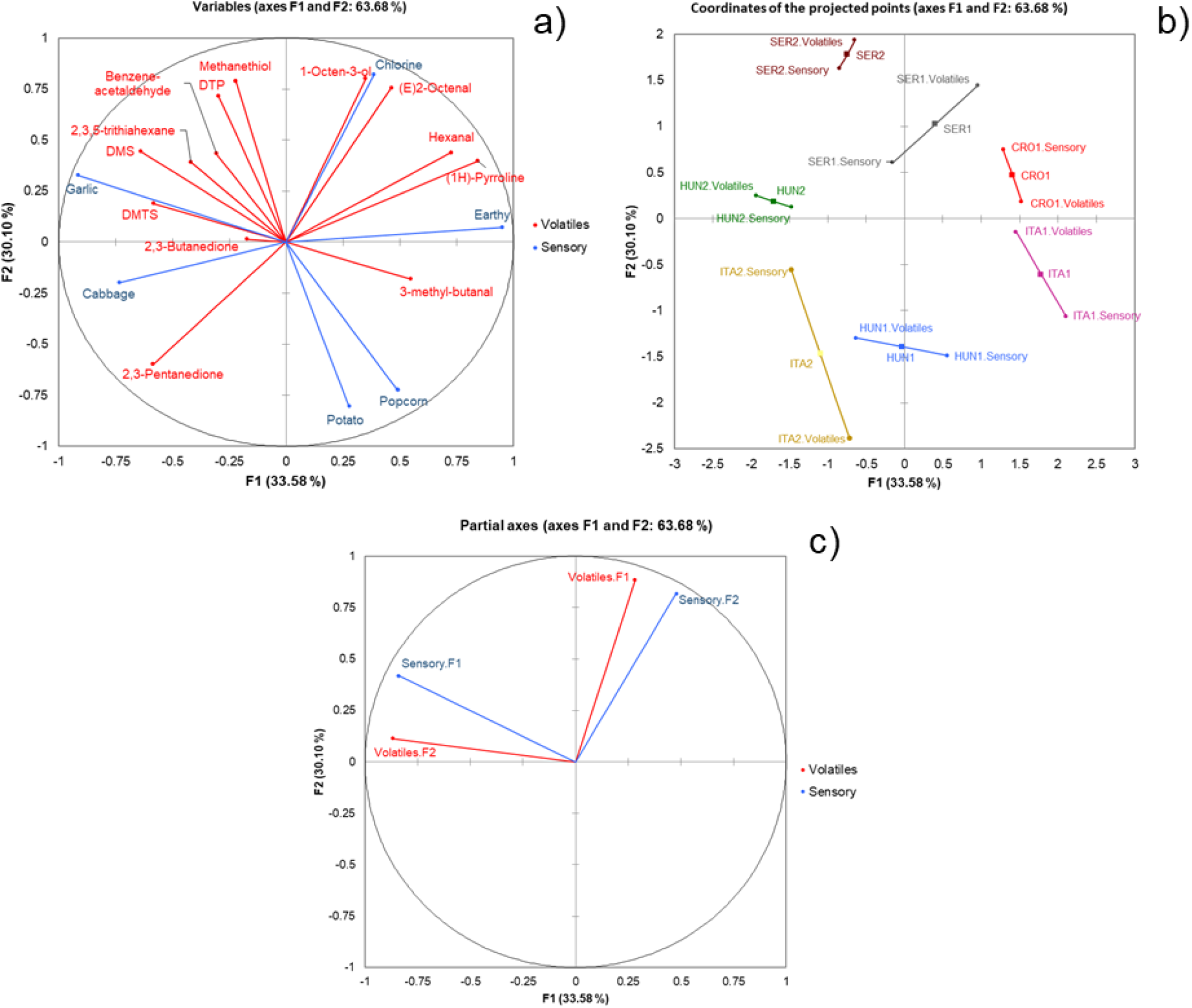
MFA plots exploring the correlations between odour active volatile compounds of pooled fruiting bodies by site and the sensory properties determined by RATA of the truffle fruiting body extracts. a). loadings, b). scores, and c) partial axes of the two data sets for the first two components. The overall explained variances of the first two factors accounted for 63.7% of the variation of both volatiles and sensory data together. The RV coefficient determined between the two data sets were acceptable at 0.631. The direction of the partial axes for the first two components of the volatiles and sensory data sets showed good agreement in the discrimination of scores.

### Volatile profiles of individual fruiting bodies are not explained by maturity or provenance

In a next step, we aimed at identifying factors that could potentially explain variability in volatile profiles among white truffle samples, in particular the biogeographic origin and the level of maturity. To do so, we acquired the full GC-MS volatile profiles, including odour active and non-active of all fruiting bodies collected in the 7 sites of the four regions. A total of 53 compounds were detected in the headspace of individual fruiting bodies. Most of the fruiting bodies were characterised by high levels of DTP and DMS, especially with Serbian truffles (Table **S2**). PCA of the volatile profiles did not highlight a discrimination of fruiting bodies based on maturity nor provenance (Fig 5). This observation was supported by PERMANOVA analysis: neither site nor maturity were significant factors on the total variability of volatile profiles of individual truffle fruiting bodies (Bray-Curtis distance, *P = 0*.*411* and *P = 0*.*159*, respectively). Further, no significant site × maturity interaction effect was found (Bray-Curtis distance, *P = 0*.*102*). Instead, strong variations of profiles were observed between fruiting bodies collected from the same site or with the same maturity level. Samples from SER1 and 2 were the exception, where the fruiting body variation was minimal in comparison to the fruiting bodies from the other sites. In comparing the average standard deviations of across all 53 volatile compounds for each site (CRO1 (0.0029), HUN1 (0.0029), HUN2 (0.0057), ITA1 (0.0139), ITA2 (0.0098), SER1 (0.0009), and SER2 (0.001)), it was clear that both ITA samples showed the largest variations across the individual fruiting bodies. Three samples collected in Croatia and Italy (ITA1-FB1, CRO1-FB3, and ITA2-FB1) showed very peculiar volatile profiles and drove most of the variability explained by the first axis of the PCA.

**Fig. 5.**
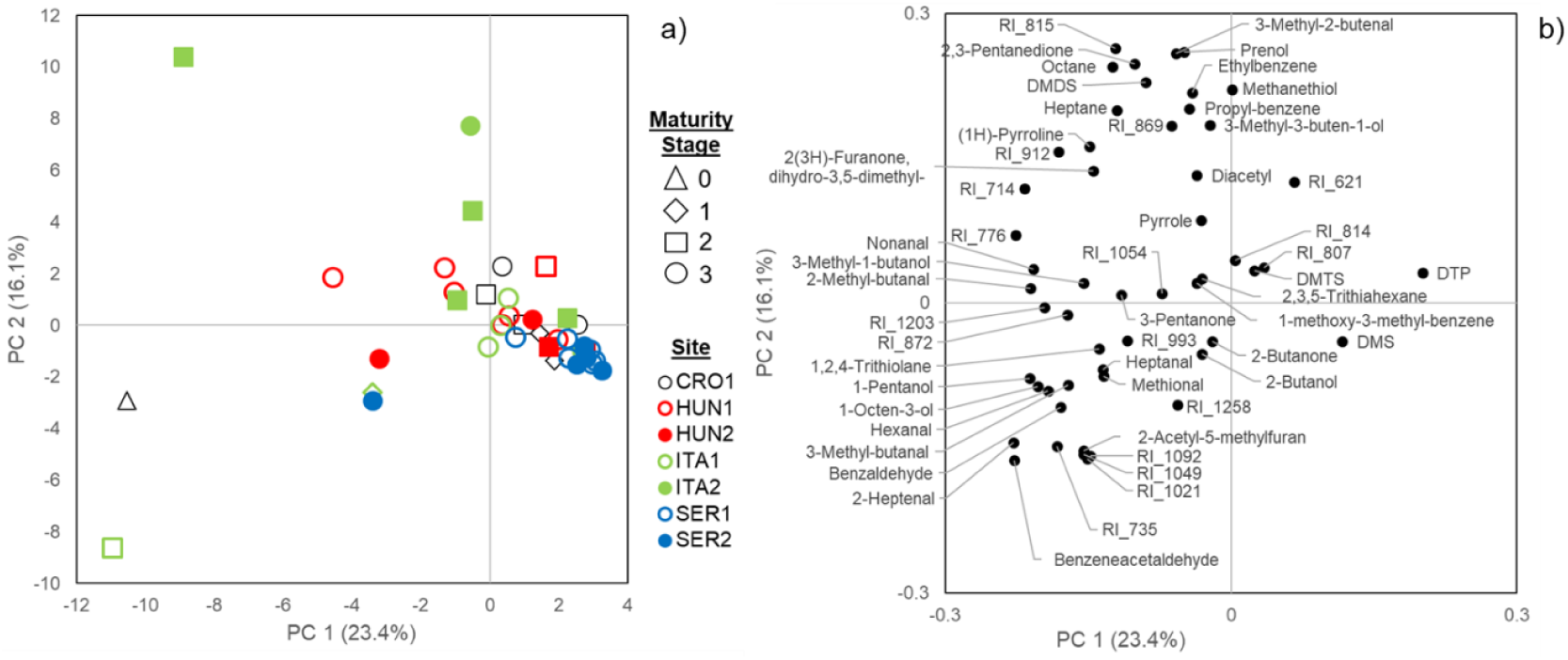
PCA scores (a) and loadings (b) plots of individual fruiting bodies and significantly different (*P <* 0.05) volatile compounds. DMS (dimethyl sulfide), DMDS (dimethyl disulfide), DMTS (dimethyl trisulfide), DTP (2,4-dithiapentane), RI (retention index; denotes for unidentified). Fruiting body maturities were categorised as percentage of asci-containing mature spores; stages 0 (0-5%), 1 (6-30%), 2 (31-70%), and 3 (70% +) (Zeppa *et al*., 2004).

Despite the overall lack of discrimination between sites based on the general profiles of volatiles, the total ion count (TIC) of 19 single volatile compounds significantly varied across sites (One way ANOVA, P<0.05). Among the 19 significantly different volatile compounds, four were previously detected with an odour through GC-O and 15 were odourless/unperceivable (Fig **S2**). 2-Butanol was only present in ITA1 (Fig **S3**), and ethyl benzene and propyl-benzene were barely detected in SER samples. Volatile compounds that were distinctly present/absent in specific sites or countries were rare, suggesting that most compounds even at the fruiting body level were ubiquitous, as already suggested by GC-O data.

### Is aroma variability explained by microbes within truffle fruiting bodies

We next investigated the composition of the microbiome of *T. magnatum* fruiting bodies and its potential influence on volatile profiles. Qualitative details of OTU diversity and abundance across the individual fruiting body samples are described in Supporting Information Note S1. Structures of the bacterial communities discriminated depending on fruiting body provenance (Fig. **6a**) but not on maturity (data not shown). Despite the variability among samples from the same site, PERMANOVA analysis indicated that sampling origin could significantly explain 26.1 % of the variability in the structures of the communities (Bray-Curtis distance, *P =* 0.009). Upon analysing the OTUs that were specific to geographical regions (Serbia, Italy, Hungary, Croatia) or collection sites, 24 OTUs (15% of all OTUs detected) and only three genera (*Chryseolina, Advenella* and *Solibacillus*) were found in a single region and all except one were found in a single site (Fig. **6b**, **Table S3**).

**Fig. 6.**
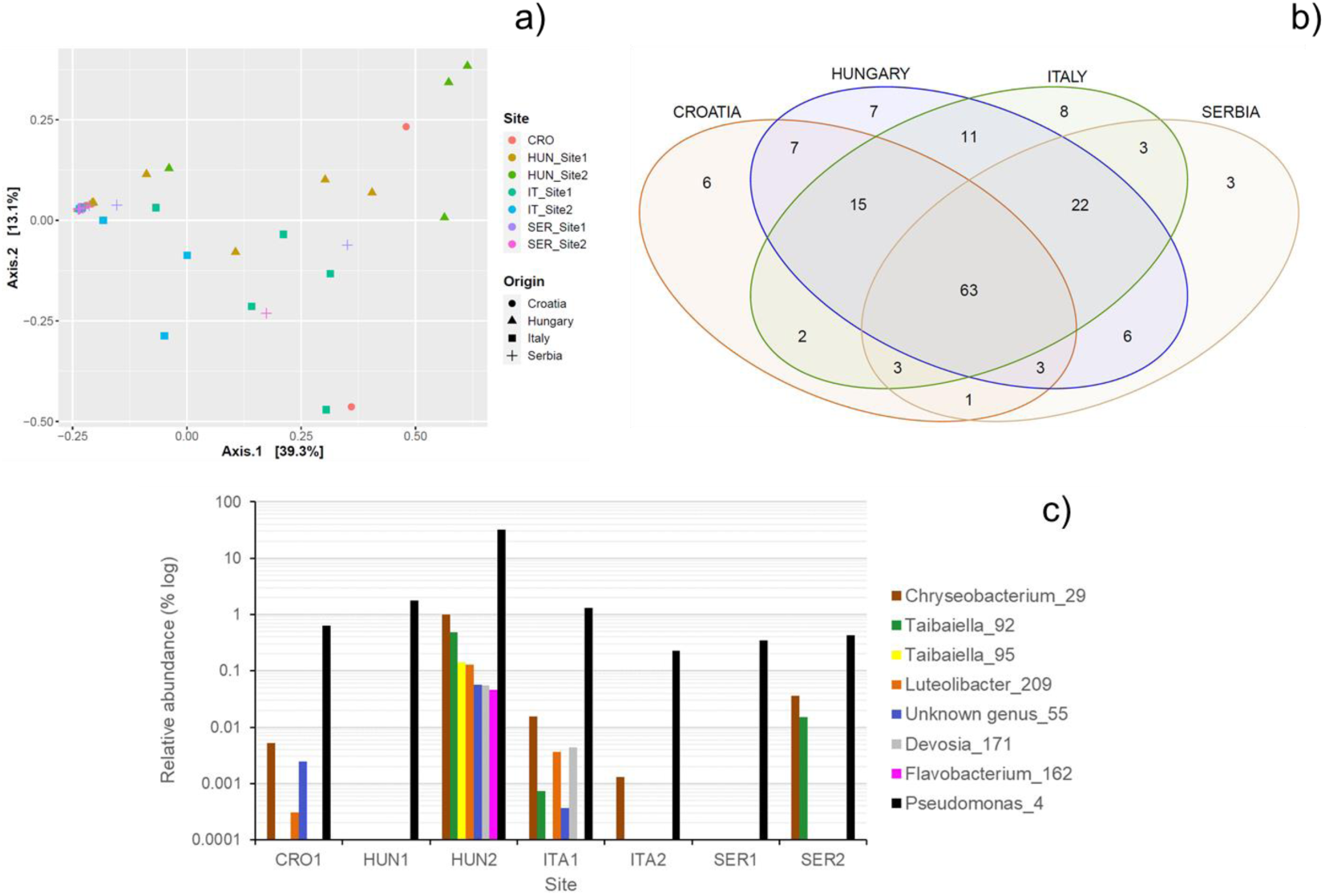
Effect of geographical origin on *T. magnatum* microbiome. a) PCoA representation of fruiting body microbial communities across sites and biogeographic area based on Bray-Curtis dissimilarity matrix, b) Venn diagram, and c) Distribution of average relative abundance of OTUs significantly enriched in Hungary site 2 compared to other sites (log scale).

These OTUs were however very rare (relative abundance being mostly < 0.02%) and they were not retrieved in all fruiting bodies collected in the area (**Table S3**). The only exceptions were the OTU 42 belonging to the genus *Chryseolina* (Bacteroidetes) that reached 12.8% of relative abundance in a fruiting body collected in Italy and *Chitinophaga* OTU 72 that was found in two fruiting bodies in Serbian Site 2 (**Table S3**). In addition, the relative abundance of 9 OTUs belonging to 8 different genera significantly varied across sites (F-test, *P –* adjust < 0.05). All 9 OTUs were significantly more abundant in fruiting bodies from HUN2 compared to other sites (Fig. **6c**). A striking difference was the massive colonization by *Pseudomonas* OTU4 of all fruiting bodies collected from HUN2. Relative abundance of this OTU varied between 8% and 79% in these fruiting bodies, while it was found at an average of 0.9% and rarely exceeded 2% in other fruiting bodies collected across the 6 other sites.

To determine any potential correlations between VOCs and microbial community composition of fruiting bodies, data was analysed with rCCA and loadings projected with covariance values greater than 0.5 are reported. Many of the fruiting bodies did not discriminate from each other, nor were there any differentiation by provenance (Fig. **7a** and **b**). Five fruiting bodies were clearly separated from others in this analysis on the consensus plot; HUN2 (three), ITA1 and SER2 (one each). It is noteworthy that microbiota of none of the five samples were dominated by *Bradyrhizobium*. Instead, microbiota had on average a higher richness and had more balanced diversity patterns than other samples (*P <* 0.01 Kruskal Wallis, **Table S4**). Several OTUs and five volatile compounds covaried closely across fruiting bodies (Fig. **7c** and **d**). Relative abundance of these OTUs was up to 4000× more abundant in some samples compared to the average in other fruiting bodies. However, their relative abundance remained below 0.1% in all cases. The presence of OTUs of the genera *Luteolibacter, Taibaiella, Devosia* and *Bosea* were associated with compound RI 814 and was specific of samples HUN2FB (fruiting body) 1 and 3. Similarly, the increased relative abundance of an OTU of the genus Allorhizobium/Neorhizobium/Pararhizobium/Rhizobium (ANPR OTU 43) and of the OTU 62 of the *Burkholderiaceae* family was associated with 2-acetyl-5-methylfuran in ITA1-FB1. The presence of these OTUs in contrast resulted in a negative covariance with elevated concentrations of methanethiol and DTP. We were not able to detect associations between the presence of specific microorganisms and the volatile compounds that are responsible for the specificities of odour active compounds through this approach. Despite this, the approach had uncovered non-odour active compounds (or compounds below their detection threshold by human nose) that associated with bacterial OTUs.

**Fig. 7.**
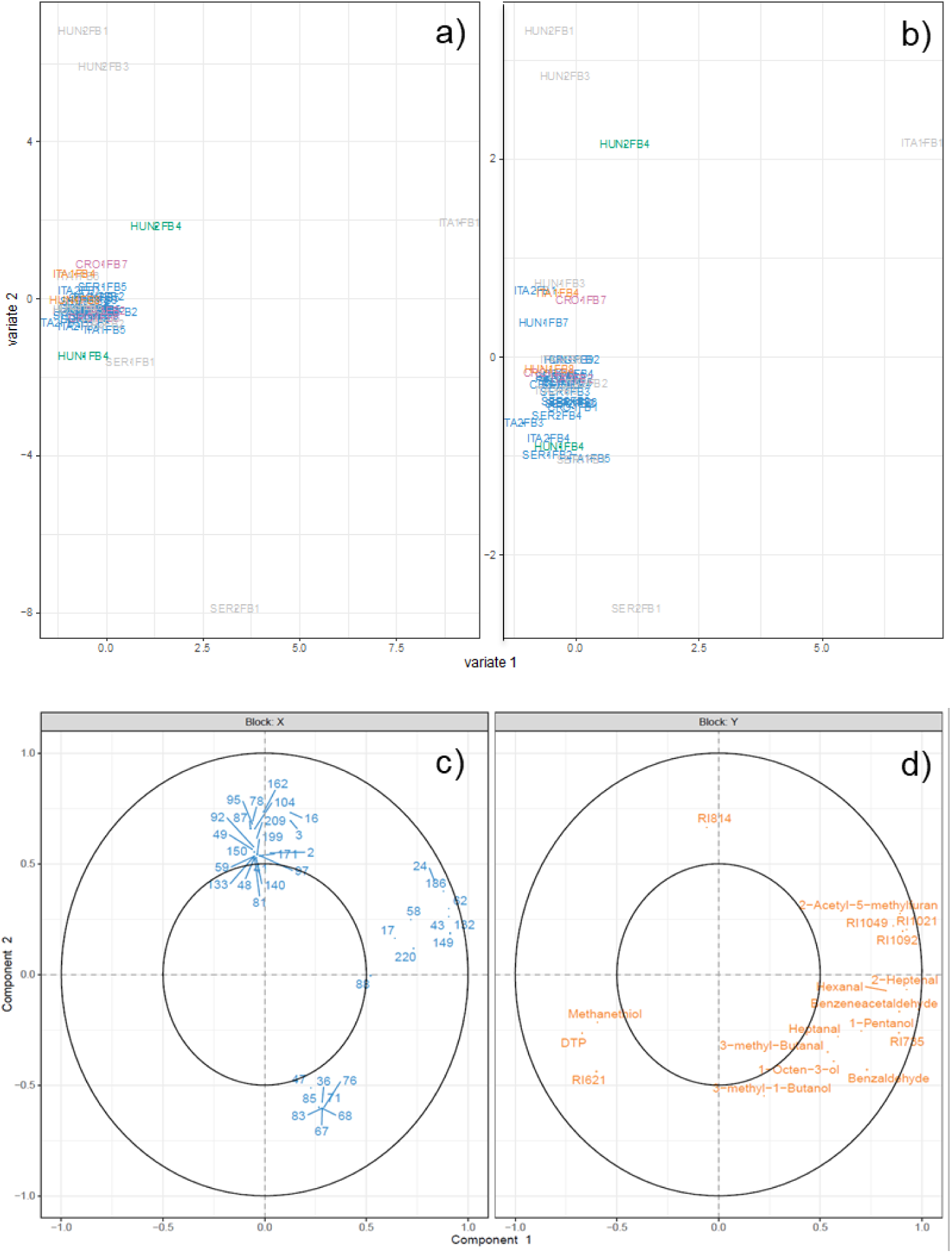
Analyses of volatile and microbiome data using rCCA. Discrimination of fruiting body scores based on a) microbiome and b) volatile profiles according to the first two variates. Correlation plot of loadings for c) bacterial OTUs and d) volatile compounds with covariance values > 0.5.

## Discussion

It has not been until recently that differences in the volatile compounds that may indicate geographical differences within *T. magnatum* white truffles have been suggested (Strojnik *et al*., 2020). There is potential for volatile differences by geographical origin to translate to perceivable differences in truffle aroma, but this had not been further investigated. In addition, the factors that are suspected to influence the volatile profile of *T. magnatum* and consequently the aroma, such as maturity and bacterial communities of fruiting bodies, had yet to be determined. The current study set out to investigate the underlying volatile compounds that may result in potential differences in the aroma of white truffle *T. magnatum* by means of GC-MS-O, and further confirm their variation when the aroma of the white truffles was sensorially evaluated. A wide multidisciplinary approach from microbiome and fungal spore morphology to volatile chemistry to sensory perception was taken, which had not been previously attempted until now.

### Odour active compounds and sensory characteristics of fresh T. magnatum

A number of volatile compounds were detected with odour characteristics, which were comparable with literature. Notably, the odour active compounds with the highest odour activity values reported previously in *T. magnatum*; DTP, 3-methyl butanal, (1H)-pyrroline, 2-methylbutanal, 2,3-butanedione, DMS, DMTS, and 1-octen-3-ol (Schmidberger & Schieberle, 2017) were also detected in the samples of the current study. A few compounds could be additionally suspected of their identity, inferred based on odour description and matching RI; RI 924 (2-acetyl-1-pyrroline) (Schmidberger & Schieberle, 2017) and RI 1181 (2-methylisoborneol) (Mahmoud & Buettner, 2016; Mahmoud & Buettner, 2017). The compound RI 807 was perceived with moderate intensity by the assessors, despite lack of identification, is suspected to be a sulfur compound based on aroma character, but further conformation is required. Curiously, very few terpenoid compounds were detected by the MS or by olfaction in any of the fruiting bodies, in agreement with Schmidberger and Schieberle (2017) but in contrast with other literature (Gioacchini *et al*., 2008; Vita *et al*., 2015; Vita *et al*., 2018). The causes of such disparity in these results is currently unknown but may lie in how the fruiting bodies were sampled for analysis. The current study had taken samples from within the gleba and not from the peridium, to reduce the measurement of volatile compounds unrelated to truffles. Unfortunately, the literature published on the investigation of white truffles rarely described sample preparation to this extent. Terpenoids nevertheless was not an important aspect of the volatile profile of *T. magnatum*, in particular from an olfaction perspective, and *T. magnatum* might lack key genes for their synthesis (Murat *et al*., 2018).

At a global level, the drivers of sensory characteristics as measured by RATA were determined through the quantified volatile data using the multiblock data analysis method MFA. This revealed that sensory characteristics specific to geographical origin were driven by concentration differences in volatile compounds, rather than presence/absence thereof. Thus, the concentration balance of volatile compound concentrations was what drove the aroma differences of truffles that originate from different sites at a global level. The comparisons of the data sets in the current study are unique in that the samples were from different matrices (fruiting bodies vs. silicon oil extracts). Ideally, the RATA data would be collected using fresh truffle fruiting bodies for a direct comparison with the volatile data.

Despite the differences in physicochemical matrices, the RV coefficient of 0.631 suggests yet a good agreement between the volatile and sensory data and indicated that the use of oil extracts for sensory assessment was a compromise that gave acceptable correlation. The wide variability seen across individual fruiting bodies within a single site, however suggests that the apparent discrimination by geographical origin from the MFA plots should be taken with caution, for reasons that will be described below.

### Variation of fruiting body volatile profiles are not determined by provenance and maturity

The influence of geographical origin on the volatile profiles of truffles have been suggested previously (Díaz *et al*., 2003; Gioacchini *et al*., 2008). The current study attempted to sample truffles from a wide geographical area spanning between southern and eastern Europe. Apart from an apparent similarity in volatile compounds of individual fruiting bodies from Serbia, differences by country of origin were unclear between samples from Italy, Croatia, and Hungary. The large average variations seen for ITA1 fruiting bodies compared to those from SER which had more than 10 fold less average variation, is likely to have an impact sensory perception. Interestingly, despite the two sites from Italy being geographically very close to each other, individual fruiting bodies from this country varied the most. Our findings contrasted literature, where fruiting bodies were able to be clustered based on volatile profiles by geographical origin in both within and across countries (Vita *et al*., 2018; Strojnik *et al*., 2020). At least at the level of orchards, volatile similarities were found in black truffle *T. aestivum* within an orchard rather than across orchards in one country (Splivallo et al., 2012). It is important to note that although our findings led us to conclude that consumers could discriminate the aroma of truffles by geographical origin, it may only take a single “rogue” truffle fruiting body that has a strong aroma characteristic from within a single site to influence the global aroma of pooled truffle fruiting bodies. That is, pooling truffle fruiting bodies essentially creates an “averaging effect” of the volatiles from one particular site.

While this addresses the practicality of sensory studies, with such high variation of volatile compounds across individual fruiting bodies, a compound could become a marker for a site simply because it is detected at high concentrations in a minority of fruiting bodies. This may have occurred for 2,3-pentanedione in ITA2 and 3-methylbutanal for ITA1, where concentrations in pooled fruiting bodies were comparatively higher across sites (Table 2), while only single fruiting bodies from their respective sites were unusually high in these two compounds.

Maturation of truffles is a complex process, where changes take place within the fruiting body ascocarps. The changes that take place are not so clear cut depending on what is being measurement about the fruiting bodies. Fruiting body weight for example has been shown to be independent of maturity for *T. aestivum* (Büntgen *et al*., 2017). Meanwhile, composition of fruiting bodies has been reported to change with fruiting body maturation, such as increases in monosaccharides and select free amino acids found in *T. melanosporum* (Harki et al., 2006) or the reduction in total phenolic content and tannins in *T. aestivum* (Shah *et al*., 2020). The most common measurement for maturity however is truffle spore melanisation, where the proportion of spores within the ascocarp that develop pigments and ornamentation increases with maturity (Zeppa *et al*., 2002). The effect of maturity on changes in volatile profiles have also been previously suggested for *T. borchii*. Key compounds that corresponded to specific maturity stages of *T. borchii* were reported and hypothesised to be derived from fatty acid metabolism as well as isoprenoid biosynthesis (Zeppa *et al*., 2004).

In the current study, maturity bore no relationship with volatile profiles or microbiome in *T. magnatum*, thus were not dependent on each other. From this perspective, our findings were consistent with the lack of correlation reported in *T. aestivum* fruiting bodies (Splivallo *et al*., 2012; Molinier *et al*., 2015). Given that the current study did not balance the sampling design to cover a wide range of maturities within each sampling site, it is unknown whether the development of volatile compounds with maturity can be site dependent.

### Microbiome covaried with volatile compounds for a few selected samples

Bacterial community composition of *T. magnatum* was highly variable among fruiting bodies, no matter their origin. Like in other truffle species, members of the *Bradyrhizobium* genus dominated in many fruiting bodies (Benucci & Bonito 2016, Antony-Babu et al. 2014, Splivallo et al. 2019) but they were replaced by other bacterial taxa in 40 % of the fruiting bodies. Similar patterns of variations were found in fruiting bodies of *T. aestivum* (Splivallo et al. 2019) and *T. melanosporum* (Deveau et al. unpublished data). However, it is noteworthy that non-*Bradyrhizobium* dominant genera differed between *T. aestivum* and *T. magnatum*.

OTUs of the genus *Pedobacter*, which were dominant in the bacterial communities of fruiting bodies of certain *T. aestivum*, were found in *T. magnatum* but never dominated. Despite an overall low contribution of the biogeographic origin of truffle fruiting bodies on the structure of the bacterial communities (24% of variability), this work revealed the potential existence of bacterial markers of *T. magnatum* origin. Indeed truffles from the Hungarian site 2 were colonized by an OTU of *Pseudomonas* that was rare in truffles from all other sites. Further analysis will be required to determine whether it is a peculiarity due to the specific conditions of this site or a more generic phenomenon that can be used to track the origin of truffles on the market.

Since many bacteria have the ability to produce compounds involved in the formation of truffle volatiles (Vahdatzadeh et al. 2016), and both truffle microbiota and volatiles vary among fruiting bodies, it is tempting to speculate that there is a link between the two. To investigate this point, covariance of the microbiota and volatile compounds of the individual fruiting bodies were determined. The majority of the covariances were explained by the extreme samples characterized by unusual volatile profiles. With ITA-FB1, it was not surprising that the sample was explained by 2-acetyl-5-methylfuran due to its unusually high abundance in that fruiting body and was an anomalous sample, along with four other unidentified volatile compounds. The close covariance of several OTUs with 2-acetyl-5-methylfuran could suggest that the compound was either derived from bacteria or that some bacteria were favoured by the production of this compound. Several ascomycetes fungi can produce this compound (Ting *et al*., 2010) but nothing is known regarding the ability of bacteria to produce it and its effect on their growth so far. Although this compound has been described to have a nutty aroma (Burdock, 2010), it’s level may have been too low to have a role in the aroma of sample ITA1-FB1, as none of the assessors detected the compound through GC-O. Three samples from HUN2 site showed high covariance between an unidentified volatile compound (RI_814) with several bacterial OTUs. Further research is needed to determine these specific volatiles have an effect on the development of microbial communities or/and if they are produced by some truffle associated microorganisms.

Nevertheless, our analysis did not show any consistent discrimination of the interaction between bacteria and volatiles by provenance of truffle fruiting bodies. This implied that there was no obvious covariance between microbiota and volatile profile of truffle fruiting bodies of *T. magnatum*, that were unique to country of origin. Instead, the variation of truffle fruiting bodies as individuals drove the discrimination of the model and bacteria cannot be used as a predictor of volatile and likely aroma.

The odour active volatile compounds identified in the fruiting bodies through olfactometry did not show any strong correlations with bacterial OTUs measured in the current study. On the one hand it could be concluded that bacteria in *T. magnatum* were not responsible for the synthesis of key odour active volatile compounds, which is in contrast with findings in *T. borchii* (Splivallo *et al*., 2015). But on the other hand it is possible that many microbes could produce the same volatile compounds to the extent that extracting correlations for such information is difficult. This phenomenon has also been demonstrated for thiophene-containing compounds in *T. borchii* (Splivallo et al., 2015). Further, pure cultures of truffle mycelium are also known to produce some of the volatile compounds often reported in truffle fruiting bodies including some within the current study, such as 1-octen-3-ol and 3-methyl butanal (Vahdatzadeh & Splivallo, 2018). It is however important to mention that previous studies have attempted to characterise the production of volatile compounds by specific microbial species in vitro (Buzzini *et al*., 2005; Vahdatzadeh & Splivallo, 2018). The results of the current study reveals that the realities of volatile synthesis by microbes within truffle fruiting bodies in nature may be more complex than once thought. With low covariations between odour active volatile compounds and bacteria, other factors may possibly control the synthesis and hence the variability of volatile compounds within the fruiting bodies, such as genetic diversity as found in *T. aestivum* fruiting bodies (Molinier *et al*., 2015). Genetic variability has been reported for *T. magnatum* (Mello *et al*., 2005) and further separated into groups based on genetic structure (Belfiori *et al*., 2020), but its extended influence on volatile profile variation for this truffle species is still yet to be determined. One aspect that is noteworthy of mention, is that the microbiome measurements focused only on bacterial communities and any fungal species that may have been present in the fruiting bodies were not measured. Although yeasts have been mainly isolated from truffle fruiting body surfaces (peridium) and not from within (gleba) (Rivera *et al*., 2010), it does not rule out the possibility of volatile compound migration from the surface to within the truffle fruiting body. Specific interactions between yeast and bacterial species are also known to result in the synthesis of volatile compounds (Frey-Klett *et al*., 2011). Important yeast species that could play a role in such synthesis of important volatile compounds from within the fruiting bodies could possibly have been missed in the current study.

### What could be the drivers of volatile/aroma and microbial variation?

Fruiting bodies were measured in as “fresh” state as possible within 3 days of harvesting. It is important to note that a reduced time from harvest to measurement does not necessarily indicate absolute fruiting body freshness. It is possible for the fruiting bodies to begin deteriorating within the soil, which will undoubtedly affect all measures made in the current study. All samples were arbitrarily checked for firmness before fruiting body processing, which is an anecdotal indicator of freshness. It is possible that changes may already be taking place within the fruiting body while the firmness remains unchanged. A follow up study is required to monitor volatile and microbial changes in *T. magnatum* fruiting bodies with storage time.

More pressing however, is the fact that truffle availability is season and weather dependent, making harvest unpredictable. Given that *T. magnatum* is notoriously difficult to cultivate in orchards (Riccioni *et al*., 2016), variability of volatiles and microbiome in the fruiting bodies are dependent on nature. There was an inherently large amount of variation in volatile and microbiome data between the fruiting bodies of the truffles even within a single site. It is possible that host trees in association with the truffles was a source of the wide variation seen in the current study, as previously suggested for *T. melanosporum* (Culleré *et al*., 2017). In the current study, the host trees were not provided by truffle hunters, as to the best of our knowledge, *T. magnatum* fruiting bodies were all wild, which may have added to variability. Other factors that may play a role in fruiting body variation may be related to soil characteristics, environmental input such as water availability and microclimate weather patterns of the local areas.

To conclude, this study was the first to explore underlying relationships between fruiting body maturity, microbiome, volatile profiles, and sensory perception of *T. magnatum* fruiting bodies. Maturity did not play a role in the variation of volatile profiles or bacterial communities in the fruiting bodies. The variation in truffle aroma were discernible by chemical means as well as by olfaction of single compounds. This extended to perceptual discrimination of truffle extracts from global aroma assessment at the level of consumers.

Key aroma compounds that were ubiquitous across all truffles, drove the differences in aroma perception across sampling sites. Consistent discrimination of fruiting bodies through volatile compounds and microbiome as a function of provenance were not found. However several key bacterial species have suggested a close relationship with key volatile compounds, albeit non-odour active and the covariances tended to explain only extreme samples. The contribution of bacterial community within fruiting bodies on volatile profiles of *T. magnatum* truffles remains unclear and thus the underlying drivers of aroma variation of the truffle fruiting bodies are yet to be determined. The findings in the current study are a first step in paving the way for further investigations to determine in detail the relationships between bacterial species, volatile profiles, and aroma perception.

## Acknowledgements

The authors would like to acknowledge all the participants who tirelessly smelt the samples during the olfactometry trial. JN acknowledges the consumers who voluntarily participated in the RATA study. The authors would like to thank the truffle hunters in complying with the scientific logistics of the study. JN’s research was supported by the Alexander von Humboldt Foundation. The study was approved by the ethics committee of Goethe University. The Deutsche Forschungsgemeinschaft (DFG) is acknowledged for financing the installation of GC-MS peripherals. RS and JN thank Žaklina S. Marjanović for kindly providing truffle fruiting body samples. AD was supported by the French National Research Agency through the Laboratory of Excellence ARBRE (ANR-11-LABX 0002 01).

## Author contribution

JN and RS contributed to the conception of the experimental concept. JN, RS, and AD contributed to the data collection, analysis, interpretation, and preparation of the manuscript.

## Conflict of interest

JN and AD do not declare any conflict of interests. RS is involved in a start-up company Nectariss Sàrl.

## Notes

### Competing Interest Statement

JN and AD do not declare any conflict of interests. RS is involved in a startup company Nectariss Sarl commercializing truffle flavors.

