## Supplemental information for "Geographical based variations in white truffle *Tuber magnatum* truffle aroma is explained by quantitative differences in key volatile compounds"

Article title: **The mystery of *Tuber magnatum* truffle aroma continues: aroma does not consistently vary by country of origin, fruiting body maturity, or bacterial community.**

Authors: **Jun Niimi, Aurélie Deveau, Richard Splivallo**

The following Supporting Information is available for this article:

**Methods S1** ***Truffle fruiting body mass and maturities***

**Fig. S1** Box and whisker plots of individual fruiting bodies from seven sites; a) mass and b) maturity. Superscripts above the box and whiskers of maturities by region denote for significantly different means using Tukey's honest significant different post hoc test.

**Methods S2** ***Volatile analysis using Gas chromatography-mass spectrometry and olfactometry (GC-MS &O)***

**Methods S3** ***Microbiome analysis***

**Methods S4** ***Sensory evaluation of truffle extracts using rate all that apply (RATA)***

**Table S1** List of attributes with definitions derived from GC-O analyses provided to the consumers.

***Table S2*** Volatile profile of individual fruiting bodies, measured using the GC-MS.

**Notes S1** ***Bacterial communities in T. magnatum fruiting bodies***

*Methods S1 – Truffle fruiting body mass and maturities*

Truffle maturity were determined using previous methods (Zeppa et al., 2004) by estimating the percentage of ascii containing immature spores. Fruiting body maturity and masses were analysed with descriptive statistics and analysed with one-way ANOVA. The two data sets were analysed with Pearson correlation to determine the correlation between the two measurements. Single fruiting body weight ranged from 7.3 to 99.3 g and fruiting body maturities ranged from 1 to 94.5 % (Fig **S1**). The mass of fruiting bodies despite being variable, were not significantly different by region (Fig **S1a**.). The maturities of each individual fruiting bodies were however significantly different by region ($p$ = 0.006) (Fig **S2b**.).

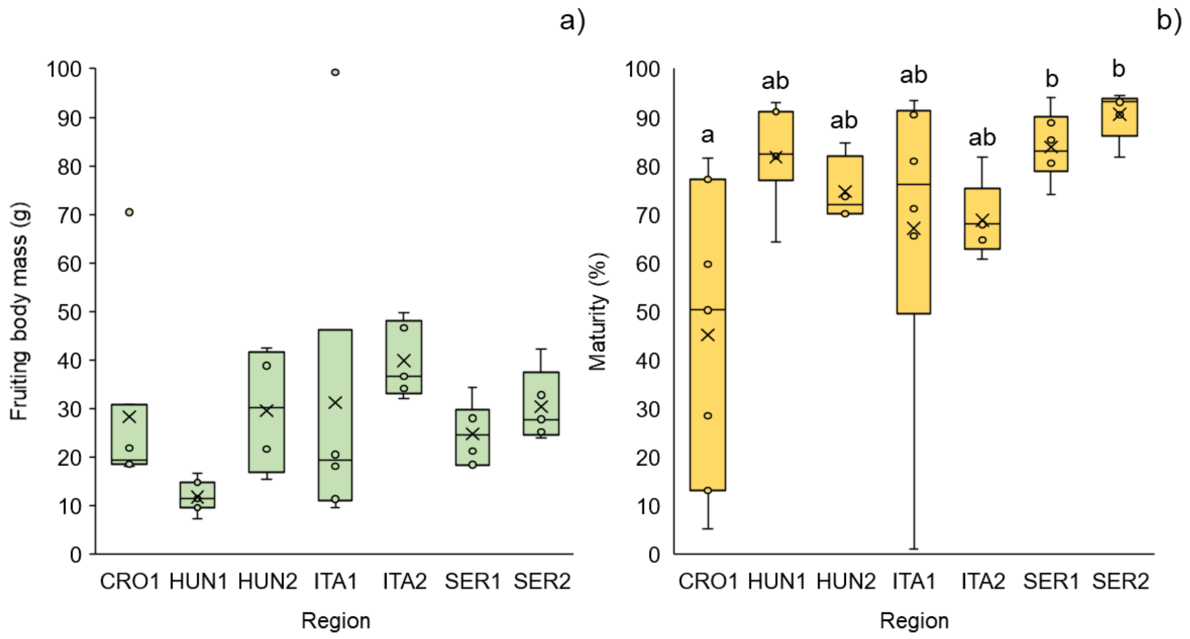

Fig **S1**. Box and whisker plots of individual fruiting bodies from seven sites; a) mass and b) maturity. Superscripts above the box and whiskers of maturities by region denote for significantly different means using Tukey's honest significant different post hoc test.

Overall, fruiting bodies from SER1 and 2, and HUN1 had the highest proportion of mature spores, while fruiting bodies from Croatia were the least mature. Within each site, large variations in the maturities of fruiting bodies were found for those from CRO1 and ITA1, which may have influenced the ANOVA. Analysis with Pearson correlation showed no significant correlation between fruiting body maturity and mass at all, suggesting that fruiting body size is independent of truffle fruiting body ripeness. This corroborated with the finding for *T. aestivum*, where fruiting body maturity bore little relationship with ripeness (Büntgen et al., 2017).

### *Methods S2 – Volatile analysis using Gas chromatography-mass spectrometry and olfactometry (GC-MS &O)*

Individual fruiting body pieces (Fig. **1-1c**) and pooled truffle fruiting bodies (Fig. **1-2b**) (300 ± 5 mg) were prepared in the SPME vials, sealed with a screw cap fitted with silicon/PFTE septum (VWR, Darmstadt). Samples were analysed for their volatile profile on the same day as the sample preparation described above. Samples were analysed using the GC (Agilent 7890B) equipped with an MS (Agilent 5877B) (Agilent technologies, Santa Clara, CA, USA) and olfactometer. The GC-MS was equipped with a PAL autosampler (CTC Analytics, Zwingen, Switzerland) with sample incubation heating/stirring chamber. Volatile compound chromatography was performed on a HP-5MS column (Agilent Technologies, Waldbronn, Germany, 30 m × 0.25 µm × 0.25 mm). Sample volatile compound extraction was performed using a 2cm fibre SPME composed of three phases with 50/30 µm thickness (DVB, CAR, PDMS) (Stableflex 23 Ga, Supelco, PA, USA). Samples were firstly equilibrated of their headspace at 60°C for 15 min, followed by extraction of volatiles at the same temperature for 30 min. The GC run program used was the same as that previously described (Vahdatzadeh & Splivallo, 2018) , but with a final hold time of 6 min at 260°C. Limit of detection was determined visually and the limit of quantification was set at 3 x the limit of detection. The MS was set to detect mass ions between m/z 40 - 350 at a scan frequency of 4.5 per sec and the detector set at a voltage of 1250V. The data was acquired using the MassHunter GCMS Data Acquisition software (v. B.07.04.2260, Agilent Technologies Inc., PA, USA) and exported as three-dimensional data (CDF).

Volatile compounds from the pooled fruiting bodies within each site that were odour active and above the limit of quantification were quantified by means of calibration curves using standards, together with the internal standard of eucalyptol (5 mg of 20 mg kg). Where standards could not be obtained for odour active compounds, their equivalence were calculated based on the standard curves of chemically similar compounds.

To determine the odour active compounds in the headspace of truffle sample, the grated fruiting body samples were used for analysis (Fig. **1-2a**). The headspace sampling and injection of the grated truffle fruiting body samples was performed according to the procedures described in 2.5.1 and analysed for odour active compounds simultaneously with MS analysis. The time temperature program was modified for the olfactometry runs, where the initial temperature of 40 °C (5 min), followed by a ramp rate of 3 °C/min till 160 °C, and finally heated to 260 °C at a rate of 75 °C/min with a final hold of 6 min. Chromatographically separated compounds were split with a four-port splitter at the end of the column (Silflow, Trajan Scientific and Medical, Australia). Compounds were split using extra helium at a ratio of 4:1 (sniff port:MS) using an ultra-inert pressfit. The split compounds were carried to sniffing port (Phaser GC olfactory port, GL Sciences B.V., Eindhoven, The Netherlands) through a transfer line (1 m). The sniffing port combined carrier gas (purified compressed air) with humidification to deliver compounds to the glass nosepiece. Four panellists previously trained in sniffing through GC-O as well as scoring intensity, participated in the olfactometry evaluation. For each sample, assessors recorded their verbal descriptions of each odour active compound together with intensity scores by clicking on a button and choose the intensity level (1, 2, and 3 as low, medium, and high intensity, respectively) on an olfactory Voicegram Recorder (v. 2.2.17, GL Sciences Inc., Tokyo, Japan). Each assessor sniffed the first 30 minutes of the GC runs, twice a day (replicates of samples), with breaks of two hours in between each sniff. Blank runs were also conducted for each assessor in replicate.

***Methods S3 – Microbiome analysis***

*DNA extraction and characterization of bacterial composition of T. magnatum fruiting bodies*

Samples were processed as described in Vahdatzadeh, Deveau, and Splivallo (2019). Briefly, after DNA extraction, quality was verified by electrophoresis and DNA was quantified with Qbit fluorometer (Thermofisher). Microbial characterization was performed using PCR-high throughput amplicon sequencing. Amplicon libraries of 16S rRNA were produced using 787r (5′-ATTAGATACCYT- GTAGTCC- 3′) (Nadkarni, Martin, Jacques, & Hunter, 2002) and 1073f (5′-ACGAGCTGACGACARCCATG -3′) primers (On, Atabay, Corry, Harrington, & Vandamme, 1998). Each primer contained a linker and a barcode which were used for the sample identification. Negative (sterile water) and positive controls were included in the library preparation. Polymerase chain reactions (PCRs) were performed in a final volume of 25 µl containing 2 µl of template DNA (10 ng.µL), 10 µl of PCR Mastermix (5 PRIME) and 1 µl of each forward and reverse primers (0.2 µM). Amplification conditions were 94 °C for 10 min, 29 cycles 94°C for 30s, 48°C for 45s, 72°C for 90s, followed by 72°C for 10 min. The concentration of PCR products was estimated by gel electrophoresis and 30µl of each amplicon was sent for MiSeq Illumina sequencing to GenoScreen (France) for 2x 250 bp Illumina Miseq sequencing.

*Sequence processing*

Bacterial raw sequences were processed with FROGS (Find Rapidly OTU with Galaxy Solution) implemented on the Galaxy analysis platform. Sequences were demultiplexed, dereplicated, sequence quality was checked, oligonucleotides, linker, pads and barcodes were removed from sequences. Then sequences were removed from data set, if they were non-barcoded, exhibited ambiguous bases or did not match expected size (286 bp). Remaining sequences were clustered into operational taxonomic units (OTUs) based on the iterative Swarm algorithm, then chimeras and phiX contaminants were removed. OTUs with a minimum number of reads above $5.10^{-5}$ percent of total abundance were kept for further analyses. Bacterial affiliation was performed by blasting OTUs against SILVA database. OTUs with blast identity < 90% and <90 % coverage were considered as potential chimera and were removed from the dataset. Finally, OTUs corresponding to chloroplasts or mitochondria were removed from the data set. Per-sample rarefaction curves were produced to assess sampling completeness, using function *rarecurve()* in package Vegan v3.5-1 (72) in R (version 3.4.3 ; 73).
Based on these, subsequent analyses of diversity and community structure were performed on datasets where samples had been rarefied with the Phyloseq package to achieve equal read numbers according to the minimum number of total reads in any sample (45,846 reads).

*Statistical analyses*

Statistical analyses and data representations were performed using R software (73, R studio v1.2.5001). Differences between bacterial community structures of fruiting bodies collected from different sites were tested using permutational multivariate analysis of variance (pairwise PERMANOVA) based on Bray-Curtis distances and differences in structures were visualized using Principal Coordinate Analysis (PcoA) using Bray-Curtis dissimilarity matrix. Fisher test followed by a Benjamini and Hochberg correction (fdr correction) was used to detect significant differences in the relative abundance of bacterial OTUs between sites. Venn diagram was produced from binarized dataset using the limma package.

***Methods S4 – Sensory evaluation of truffle extracts using rate all that apply (RATA)***

Truffle extracts prepared with silicon oil (2c., Fig. 1) and one commercial truffle oil (Bartolini, Arrone, Italy) (dummy sample) were pipetted (500 µL) into wide-neck Erlenmeyer flasks (100mL) and covered with aluminium foil, two hours prior to evaluation. Consumers recruited from Goethe University Riedberg Campus were recruited for the evaluation of the truffle extracts (n = 81). All consumers signed an informed written consent prior to the evaluation. The task evaluation procedures were introduced at the beginning of the test to the consumers whilst they were not informed of the true objective of the study. During the introduction, the assessors were provided with packing sheets that outline the basic instructions as well as the order of samples to be assessed using the online data acquisition software RedJade® (Redwood Shores, CA, USA). To assist the consumers in understanding of the aroma descriptions, a list of attributes for RATA along with their definitions, which was derived from terms generated from GC-O (Table 2). Synonymous words used during GC-O were compiled together and formed as definitions. The consumers answered a prequestionnaire consisting of their age range, gender, and familiarity of truffle products. The first sample evaluated was a commercial truffle oil to serve as a practise sample for the consumers to familiarise with the RATA procedure. With this first sample, consumers were instructed on evaluation procedures, use of scales, and the user interface of the software. Consumers smelled the aroma of the samples and evaluated the intensities of attributes on a 7-point scale. The order at which the attributes were listed on the computer screen were randomised for every sample per consumer. After the evaluation of the dummy sample, the consumers evaluated the test samples individually.

Samples were presented in randomised orders and labelled with blind three-digit codes, apart from the dummy sample which was always presented in the first position. Evaluation took place in a large open room equipped with computers, where up to 20 consumers could be tested at a time. Samples were presented at room temperature (22°C) under natural lighting. RATA testing was conducted over three days and the trial was ethically approved by the Goethe University.

**Table S1** List of attributes with definitions derived from GC-O analyses provided to the consumers.

| Attribute | Definition |
|---|---|
| Garlic | The smell of fresh garlic |
| Cheese | The smell of cheese |
| Earthy | The smell of dirt or soil |
| Mushroom | The smell of fresh mushroom |
| Potato | The smell of cooked potatoes |
| Cabbage | The smell of cabbage |
| Vegetable | The smell of vegetables |
| Butter | The smell of butter |
| Popcorn | The smell of popcorn |
| Grassy | The smell of grass/plants |
| Floral | The smell of flowers |
| Malty | The smell of malt |
| Broth | The smell of broth/soup |
| Chlorine | The smell of chlorine/sperm |

***Table S2*** Volatile profile of individual fruiting bodies, measured using the GC-MS.

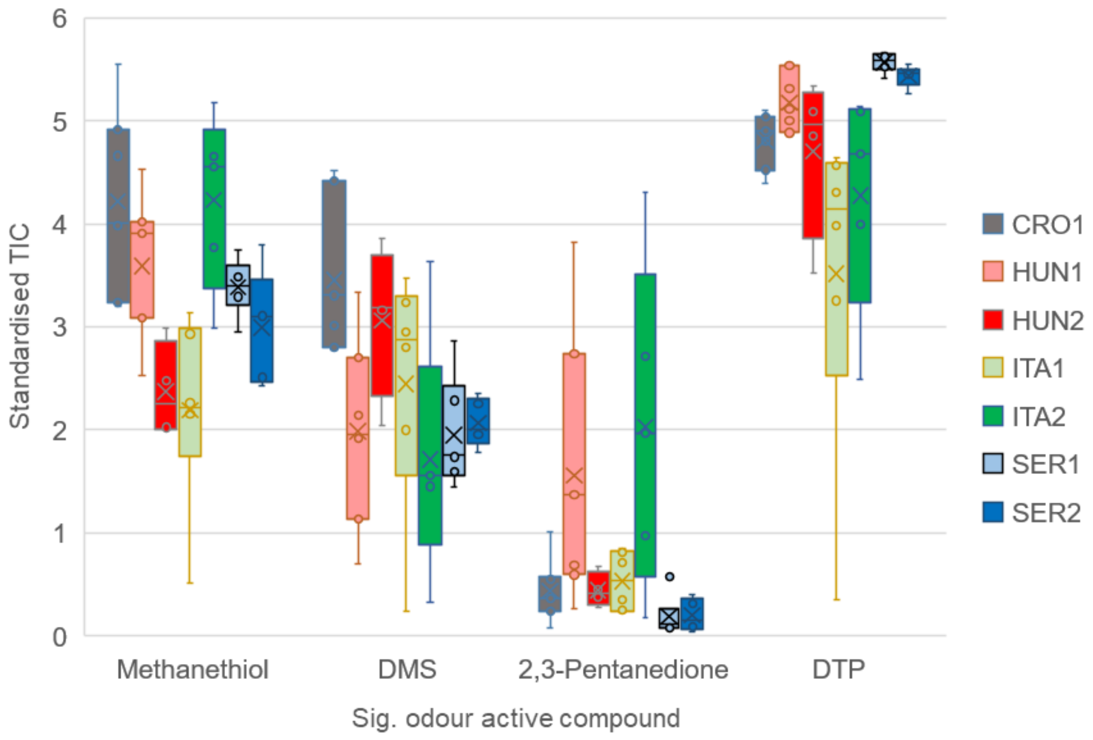

Fig **S2.** Variation of significantly different (*P* < 0.05) odour active compounds measured from individual fruiting bodies across sites. Standardisation was performed by scaling data 1/standard deviation.

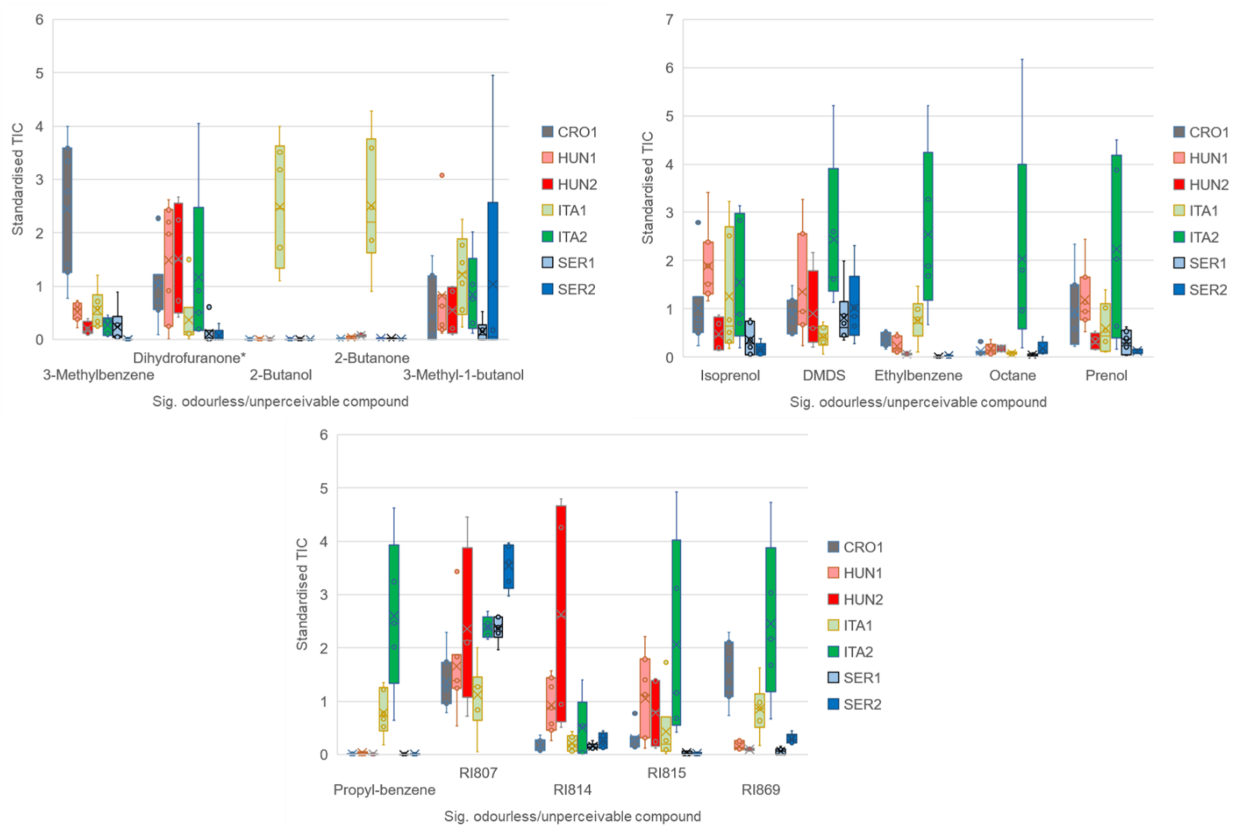

Fig **S3.** Variation of significantly different ($P < 0.05$) odourless/unperceivable compounds measured from individual fruiting bodies across sites. *Dihydro-3,5-dimethyl-2(3H)-furanone. Standardisation was performed by scaling data 1/standard deviation.

***Note S1 – Bacterial communities in T. magnatum fruiting bodies***

A total of 158 OTUs were detected among the 40 fruiting bodies analysed in this study. Fruiting bodies were colonized on average by 42 OTUs but varied depending on the fruiting bodies as it ranged between 19 and 76 OTUs per fruiting body (Fig. **S4a**). Similarly, diversity varied between samples. Although most bacterial communities were dominated by a few OTUs as indicated by low Inverse Simpson diversity index, some communities showed more balanced patterns. In accordance with this, most OTUs were rare in terms of abundance and of the number of fruiting bodies in which they were encountered (Fig. **S4b**): 40% of all OTUs were retrieved in five fruiting bodies at most and these OTUs represented no more than 1.3% of all reads. By contrast the most abundant OTU accounted for 64% of all reads. Five OTUs were found in all fruiting bodies and they were also among the most abundant (Fig. **S4b**).

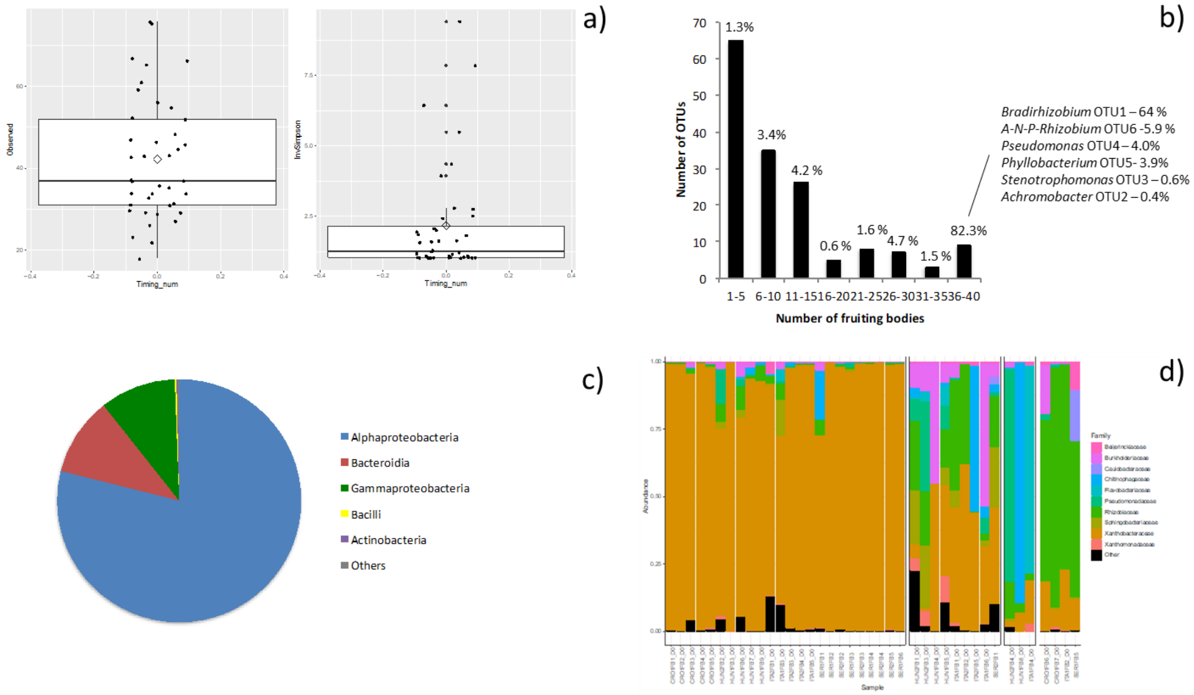

Fig. **S4** Diversity and composition of bacterial communities of *T. magnatum*.

Overall, communities were dominated by OTUs affiliated to α-Proteobacteria (78.9 % of the reads in average) followed by Bacteroidia (10.4 %), γ-Proteobacteria (10.1 %). Bacilli and Actinobacteria were rare (< 0.2%) but they were found in most fruiting bodies. Members of the Verrucomicrobiae, Clostridia and Thermoleophilia classes were also detected but at very low levels in some fruiting bodies only (Fig. **S2c**). The composition of the bacterial communities was very heterogeneous depending on the fruiting bodies. While 60% of the fruiting bodies showed communities largely by members of the Xanthobacteraceae family dominated (>70% in relative abundance) and in particular by one OTU of the *Bradyrhizobium* genus, the 16 fruiting bodies were characterized by other patterns of community composition (Fig. **S4d**), 9 of which showing patterns of several families being “co-dominant”. Other fruiting bodies were characterized by communities dominated by OTUs belonging to a single family: Rhizobiaceae, Chitinophagaceae, Flavobacteriaceae or Pseudomonadaceae. We observed in 6 cases the replacement of the dominant OTU of *Bradyrhizobium* by another single dominant OTU, either *Chitinophaga* (Bacteroides, Hun1FB8), Flavobacterium (Bacteroides, ITA1FB4), *Phyllobacterium* (α-

Proteobacteria, CRO1FB7), *Pseudomonas* (γ-Proteobacteria, HUN2FB4) or Allorhizobium group (α-Proteobacteria, ITA1FB2, CRO1FB6) (Fig. **S5**).

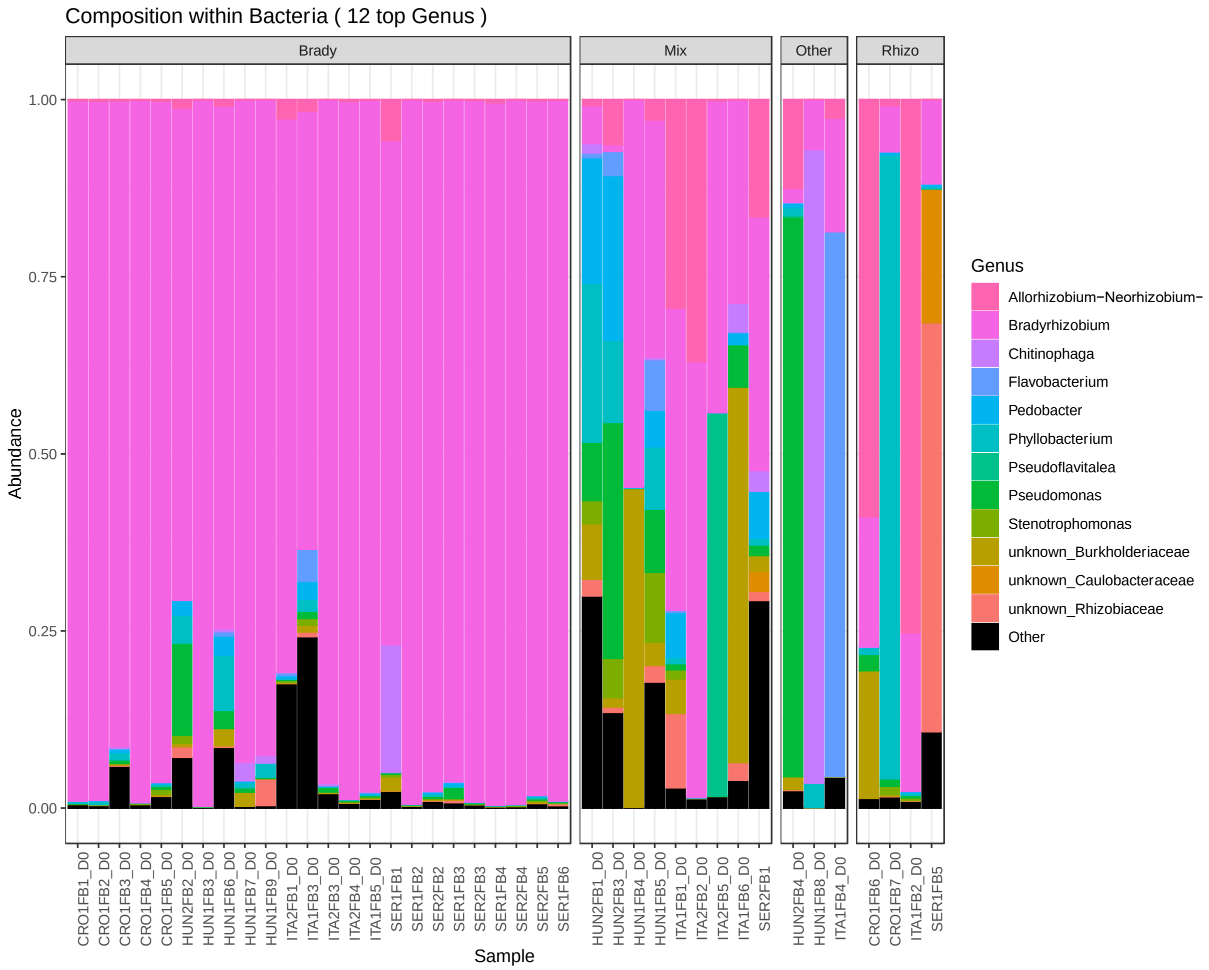

Fig. **S5** Composition of bacterial communities of *T. magnatum* at the genus level (12 most abundant genera).

| | | | | CROATIA | | | HUNGARY | | | | | | | ITALY | | | | | | SERBIA | | |
|---|---|---|---|---|---|---|---|---|---|---|---|---|---|---|---|---|---|---|---|---|---|---|
| | | | | SITE 1 | | | SITE 1 | | | SITE 2 | | | | SITE 1 | | SITE 2 | | | | SITE 1 | SITE 2 | |
| OTU ID | Class | Family | Genus | FB3 | FB6 | FB7 | FB6 | FB7 | FB9 | FB1 | FB2 | FB3 | FB4 | FB3 | FB5 | FB1 | FB2 | FB3 | FB4 | FB5 | FB1 | FB3 |
| 173 | Gammaproteobacteria | Enterobacteriaceae | unknown genus | 0,84 | 0 | 0 | 0 | 0 | | 0 | 0 | 0 | 0 | 0 | 0 | 0 | 0 | 0 | 0 | 0 | 0 | 0 |
| 69 | Actinobacteria | Micrococcaceae | unknown genus | 0,002 | 0,015 | 0 | 0 | 0 | | 0 | 0 | 0 | 0 | 0 | 0 | 0 | 0 | 0 | 0 | 0 | 0 | 0 |
| 44 | Bacilli | Paenibacillaceae | Paenibacillus | 0 | 0,009 | 0 | 0 | 0 | | 0 | 0 | 0 | 0 | 0 | 0 | 0 | 0 | 0 | 0 | 0 | 0 | 0 |
| 166 | Gammaproteobacteria | Burkholderiaceae | unknown genus | 0 | 0 | 0,04 | 0 | 0 | | 0 | 0 | 0 | 0 | 0 | 0 | 0 | 0 | 0 | 0 | 0 | 0 | 0 |
| 153 | Bacteroidia | Weeksellaceae | Chryseobacterium | 0 | 0 | 0,011 | 0 | 0 | | 0 | 0 | 0 | 0 | 0 | 0 | 0 | 0 | 0 | 0 | 0 | 0 | 0 |
| *134* | *Bacilli* | *Planococcaceae* | *Solibacillus* | 0 | 0 | 0,011 | 0 | 0 | | 0 | 0 | 0 | 0 | 0 | 0 | 0 | 0 | 0 | 0 | 0 | 0 | 0 |
| 75 | Bacilli | Carnobacteriaceae | Carnobacterium | 0 | 0 | 0 | 0,057 | 0,007 | | 0 | 0 | 0 | 0 | 0 | 0 | 0 | 0 | 0 | 0 | 0 | 0 | 0 |
| 80 | Gammaproteobacteria | Xanthomonadaceae | unknown genus | 0 | 0 | 0 | 0 | 0 | 0,002 | 0 | 0 | 0,009 | | 0 | 0 | 0 | 0 | 0 | 0 | 0 | 0 | 0 |
| 162 | Bacteroidia | Flavobacteriaceae | Flavobacterium | 0 | 0 | 0 | 0 | 0 | | 0,087 | 0 | 0,059 | 0,04 | 0 | 0 | 0 | 0 | 0 | 0 | 0 | 0 | 0 |
| 200 | Gammaproteobacteria | Burkholderiaceae | Achromobacter | 0 | 0 | 0 | 0 | 0 | | 0 | 0,002 | 0,039 | 0 | 0 | 0 | 0 | 0 | 0 | 0 | 0 | 0 | 0 |
| 52 | Bacilli | Paenibacillaceae | Paenibacillus | 0 | 0 | 0 | 0 | 0 | | 0 | 0 | 0,0065 | 0 | 0 | 0 | 0 | 0 | 0 | 0 | 0 | 0 | 0 |
| 119 | Bacteroidia | Spirosomaceae | Dyadobacter | 0 | 0 | 0 | 0 | 0 | | 0 | 0 | 0 | 0,73 | 0 | 0 | 0 | 0 | 0 | 0 | 0 | 0 | 0 |
| 246 | Bacteroidia | Flavobacteriaceae | Flavobacterium | 0 | 0 | 0 | 0 | 0 | | 0 | 0 | 0 | 0 | 0,020 | 0 | 0 | 0,002 | 0 | 0 | 0 | 0 | 0 |
| 193 | Thermoleophilia | unknown family | unknown genus | 0 | 0 | 0 | 0 | 0 | | 0 | 0 | 0 | 0 | 0,017 | 0 | 0 | 0 | 0 | 0 | 0 | 0 | 0 |
| 158 | Bacilli | Staphylococcaceae | Staphylococcus | 0 | 0 | 0 | 0 | 0 | | 0 | 0 | 0 | 0 | 0 | 0,002 | 0 | 0 | 0 | 0 | 0 | 0 | 0 |
| ***42*** | ***Bacteroidia*** | ***Microscillaceae*** | ***Chryseolinea*** | 0 | 0 | 0 | 0 | 0 | | 0 | 0 | 0 | 0 | 0 | 0 | **13,0** | 0 | 0,020 | 0 | 0 | 0 | 0 |
| 99 | Bacteroidia | Weeksellaceae | Chryseobacterium | 0 | 0 | 0 | 0 | 0 | | 0 | 0 | 0 | 0 | 0 | 0 | 0 | 0,009 | 0 | 0 | 0 | 0 | 0 |
| 168 | Alphaproteobacteria | unknown family | unknown genus | 0 | 0 | 0 | 0 | 0 | | 0 | 0 | 0 | 0 | 0 | 0 | 0 | 0 | 0,68 | 0 | 0 | 0 | 0 |
| 112 | Bacteroidia | Spirosomaceae | Dyadobacter | 0 | 0 | 0 | 0 | 0 | | 0 | 0 | 0 | 0 | 0 | 0 | 0 | 0 | 0,004 | 0 | 0 | 0 | 0 |
| *122* | *Gammaproteobacteria* | *Burkholderiaceae* | *Advenella* | 0 | 0 | 0 | 0 | 0 | | 0 | 0 | 0 | 0 | 0 | 0 | 0 | 0 | 0 | 0,092 | 0 | 0 | 0 |
| 196 | Bacteroidia | Sphingobacteriaceae | unknown genus | 0 | 0 | 0 | 0 | 0 | | 0 | 0 | 0 | 0 | 0 | 0 | 0 | 0 | 0 | 0 | 0,017 | 0 | 0 |
| **72** | **Bacteroidia** | **Chitinophagaceae** | **Chitinophaga** | 0 | 0 | 0 | 0 | 0 | | 0 | 0 | 0 | 0 | 0 | 0 | 0 | 0 | 0 | 0 | 0 | **2,79** | 0,004 |
| 86 | Bacteroidia | Spirosomaceae | Dyadobacter | 0 | 0 | 0 | 0 | 0 | | 0 | 0 | 0 | 0 | 0 | 0 | 0 | 0 | 0 | 0 | 0 | 0,039 | 0 |

**Table S3.** Relative abundance of OTUs in a single geographical area or in a single site. Relative abundances > 1% are highlighted in bold letters. Italics indicate genera that have a unique OTU in the entire dataset.

**Table S4** Bacterial richness and Inverse Simpson values of individual truffle fruiting bodies

| Sample ID | Origin | Richness | Inverse Simpson |
|---|---|---|---|
| HUN2FB1 | Hungary | 75 | 9.41 |
| HUN2FB3 | Hungary | 66 | 6.43 |
| HUN2FB4 | Hungary | 61 | 1.56 |
| HUN2FB2 | Hungary | 65 | 2.00 |
| ITA1FB1 | Italy | 67 | 3.94 |
| SER2FB1 | Serbia | 52 | 5.55 |
| Average | | 64.3 | 4.82 |
| SE | | 3.09 | 1.20 |
